# CLIMATIC NICHE DIFFERENTIATION ACCOMPANIED THE RADIATION OF LEAF-EARED MICE IN THE *PHYLLOTIS DARWINI* SPECIES GROUP (SIGMODONTINAE, CRICETIDAE)

**DOI:** 10.64898/2026.05.06.723104

**Authors:** Marcial Quiroga-Carmona, José H. Urquizo, Naim M. Bautista, Guillermo D’Elía, Jay F. Storz

## Abstract

**Aim:** to characterize the evolution of climatic niches during the diversification of the *Phyllotis darwini* species group, in order to assess the extent to which divergences involved in radiation were associated with patterns of conservatism or divergence of climatic niches, and whether the differentiation found among climatic niches correlated with species phylogenetic relationships.

**Location:** south-central Andes, surrounding lowlands, and Patagonia, South America.

**Methods:** species climatic niches were characterized by sampling contemporaneous precipitation and temperature conditions across occurrence locations and entire distributional ranges. Climatic niches were analyzed and modeled using multivariate statistics (PCA, PERMANOVA), a maximum entropy-based algorithm, and novel methods developed to explore levels of differentiation (niche overlap test) and divergence (niche divergence test) between realized and fundamental niches. Comparative phylogenetic methods were applied using a time-calibrated phylogeny and integrating climate niche data to estimate ancestral environmental niches within geographic and environmental spaces.

**Results:** comparisons revealed low levels of climatic niche overlap, both among species’ realized niches and among their fundamental niches, suggesting high levels of niche differentiation during the diversification of *Phyllotis* species. Quantifications of niche overlap further showed that observed differences among species lay primarily in the multidimensional nature of climatic niches, as unidimensional quantifications exhibited higher levels of overlap. Evolved differences among species’ climatic niches were better fitted to a Brownian motion model of evolution, but lacked phylogenetic signal and showed no significant association with species’ phylogenetic distances.

**Main conclusions:** low levels of differentiation between ancestral climatic niches suggest that the early radiation of species in the *Phyllotis darwini* species group was promoted by geographic isolation, whereas the more recent diversification of extant species was accompanied by climatic niche differentiation, possibly involving local adaptation to regional ecoclimatic changes associated with Quaternary glacial cycles. The spatial separation of sister species, the complete divergence of their climatic niches, and the lack of phylogenetic signal in niche differences suggest a scenario of diversification in which divergences were prompted by the spatial isolation, but also by the divergent selection exerted by regional climatic differences.

## 1 INTRODUCTION

The climatic niche of a given species is defined by the prevailing temperature and precipitation conditions in the species’ occupied habitat (Soberón 2007; Bonetti & Wiens 2014; Quintero & Wiens 2013). Such conditions are important determinants of the local abundance and geographical distribution of species and also shape the evolution of physiological tolerances (Soberón 2007; Colwell & Rangel 2009). Niche conservatism is defined as the tendency of species to retain their ancestral ecological niche throughout their evolutionary history, despite changes in the inhabited environmental conditions (Ackerly et al. 2006; Wiens et al. 2010). By contrast, niche divergence between species largely reflects evolutionary differentiation in physiological tolerances to a given set of environmental conditions (Kozak & Wiens 2010). Given the importance of climatic variation in determining geographic distributions and other ecological aspects of species, such as their abundances and reproductive phenology (Schluter 2001; Ackerly et al. 2006), the idea of climatic niche evolution has been widely incorporated into geographical and ecological models of speciation (Schluter 2009; Sobel et al. 2010; Wiens et al. 2010; Hua & Wiens 2013; Jezkova & Wiens 2018). Briefly, niche conservatism can configure scenarios of speciation through vicariance by confining populations to habitat patches with suitable ecoclimatic conditions and preventing their dispersal through regions with unsuitable conditions, leading populations to become reproductively isolated (Wiens 2004; Hua & Wiens 2013). Conversely, niche divergence can conduce to conspecific populations contiguously distributed, becoming highly adapted to local and different ecoclimatic conditions, causing individuals from one area to be selected against in the other, reducing gene flow between these populations, and setting conditions to divergence and eventual speciation (Rundle & Nosil 2005; Sobel et al. 2010; Hua & Wiens 2013; Jezkova & Wiens 2018).

The relevance of climatic niche evolution to the lineage diversification and shaping of the biogeographical patterns of different groups of South American rodents has been widely recognized. For example, in the family Caviidae, strong climatic niche conservatism exhibited by its lineages limited adaptation to new environments and, together with the climatic dynamics that occurred from the Late Miocene to the Pleistocene, has promoted their diversification through allopatric speciation (da Silva et al. 2020). In octodontoid rodents, niche conservatism was also identified as a key mechanism for the diversification of the family Echimyidae, whereas lineages of the family Ctenomyidae showed stronger signals of climatic niche divergence and lower niche overlap; aspects that would have favored their diversification through vicariance (da Silva et al. 2024). In the subfamily Sigmodontinae, it was found that ancient divergences were not associated with adaptive processes, whereas the cladogenetic events that gave rise to extant species were accompanied by climatic niche divergence (Reis et al. 2018). These results are consistent with the perspectives described in previous studies (e.g., Schenk et al. 2013; Parada et al. 2013, 2015), which describe that the diversification of sigmodontine was promoted by ecological opportunity and transitions to new habitats during the colonization of South American landscapes. Nevertheless, other assessments have proposed that such diversification occurred mainly through non-adaptive processes driven by geographic isolation (Alhajeri et al. 2016; Maestri et al. 2017). These discrepancies may partially stem on these previous studies are based on phylogenetic reconstructions with incomplete taxonomic sampling, and therefore, did not fully capture the entire composition within the most speciose genera within this megadiverse group (see Bangs et al. 2025). Moreover, by focusing on diversification patterns that emerge at a macroevolutionary scale, these approaches lose resolution over processes occurring at intrageneric or interspecific levels (i.e., terminal branches and tips in a phylogeny). Even so, the available evidence consistently supports the notion that the diversification of extant sigmodontine lineages was influenced by climatic dynamics occurring from the Quaternary glaciation to the present (e.g., Pardiñas et al. 2011). This highlights the need for more studies aimed at better understanding the role of climatic niche evolution in the diversification and the establishment of biogeographical patterns within the most speciose genera of Sigmodontinae.

Leaf-eared mice in the genus *Phyllotis* constitute one of the most speciose clades within the subfamily Sigmodontinae. This genus comprises 23 species (Mammal Diversity Database 2025), arranged into three natural groups commonly referred as the *andium-amicus, osilae*, and *darwini* species groups (Steppan & Ramírez 2015; Rengifo & Pacheco 2017; Teta et al. 2022). Among these, the *darwini* species group (hereafter the *darwini* group) comprises 11 species (Quiroga-Carmona et al. 2025) that are distributed in the central-southern Andes, adjoining coastal lowlands, and Patagonia (Steppan & Ramírez 2015; Figure 1). The geographic range of this clade encompasses an extensive, climatically diverse area, characterized by open arid and semiarid landscapes, including Patagonian steppes, Mediterranean scrub, the Atacama desert and Andean Puna (Olson et al. 2001). Most divergences between extant species occurred within the last 2 Mya, and there appears to have been a pulse of speciation during the mid-to late-Pleistocene (Parada et al. 2015; Quiroga-Carmona et al. 2025). The diversification of the *darwini* group occurred during a period of great geoclimatic change in South America, whose evolutionary implications have at least been evidenced in the historical demography of some populations of *P. vaccarum* and *P. xanthopygus* (see Lessa et al. 2010; Riverón 2011). In this regard, it is remarkable that most pairs of sister species within the *darwini* group have juxtaposed or partly overlapping geographic distributions along a marked environmental gradient, in which ecoclimatic conditions become rainy and cold from north to south. This pattern of spatial segregation can be considered a first indication that their diversification could have been accompanied by the divergence of their climatic niches. Nonetheless, it could also reflect a scenario in which speciation occurred under a context of vicariance where niche conservatism was one of the driving mechanisms promoting speciation. However, to date, it remains unknown how historical climatic changes in the southern cone of South America (e.g., Quaternary glaciations) have influenced the diversification and biogeographic patterns of this group of sigmodontine mice.

**Figure 1.**
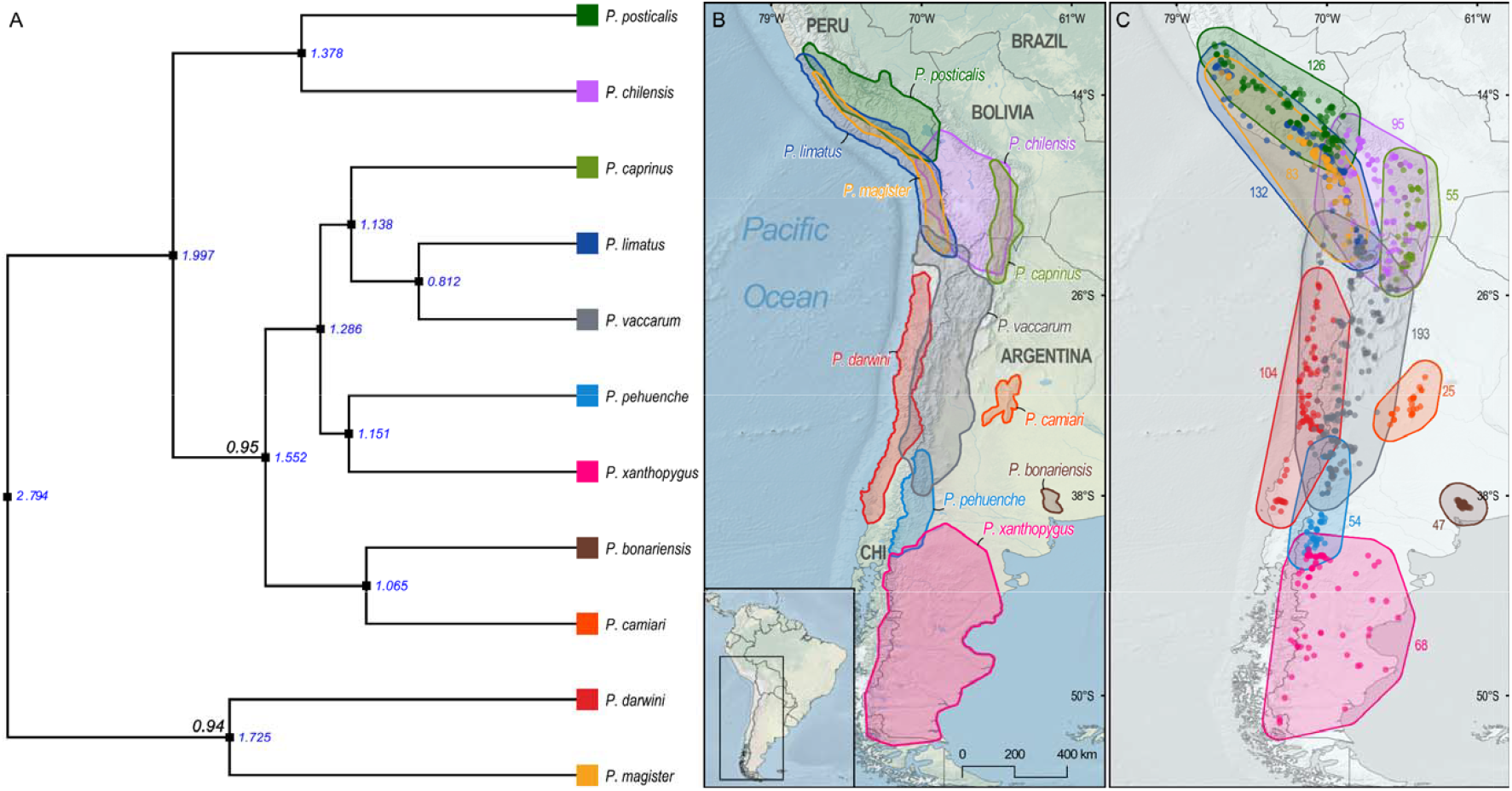
Phylogenetic relationships and geographic distributions of the Andean leaf-eared mice species within the *darwini* group. A: The pruned phylogenetic tree was obtained from the calibrated maximum clade credibility tree inferred by Quiroga-Carmona et al. (2025) using a fragment of bp of the cytochrome-b gene (Cytb). Node support is shown only for those cases in which Bayesian posterior probability values were < 1 (black numbers). The average ages of lineages splits are indicated in millions of years (blue numbers). Maps of southern South America show: B) the updated geographic distributions of Phyllotis species within the *darwini* group, according delimitations performed by Quiroga-Carmona et al. (2025), and C) the 982 presence records gathered for these species and the polygons established as their respective accessible areas or occupied distributional areas, as described in Materials & Methods. The numbers next to each polygon represent the total number of occurrences collected for each species.

Here we assess variation in climatic niches among species of the *Phyllotis darwini* group to explore which of the patterns of climatic niche evolution (conservatism or divergence) have accompanied the diversification of this clade. We used multivariate statistical approaches premised on the multidimensional conception of the ecological niches and the assumption that each species occupies its own and unique portion of the environmental niche space (Soberón 2007; Holt 2009; Peterson et al. 2011). Accordingly, the realized and the fundamental climatic niches were respectively characterized by sampling the prevailing climatic conditions present at the localities where species have been recorded, and over the areas that fully encompass their geographic ranges (Peterson et al. 2011; Guisan et al. 2017; Brown & Carnaval 2019). This information was used to quantify differences among species’ climatic niches and to develop correlative niche models (see Broennimann et al. 2012; Blonder et al. 2018; Brown & Carnaval 2019). Simultaneously, a phylogenetic approach was implemented to account for whether the differences between species’ climatic niches show a statistical association with their evolutionary relatedness by assessing the phylogenetic signal across climatic differences of species’ niches (see Wiens & Graham 2005; Losos 2008; Revell et al. 2008), and to reconstruct ancestral climatic tolerances of lineages within the *darwini* group (see Yesson & Culham 2006; Evans et al. 2009). Together, these statistical and phylogenetic approaches have provided insights into the ecogeographic scenarios (see Doebeli & Dieckmann 2004) that would have promoted the diversification of several animal groups (Knouft et al. 2006; McCormack et al. 2009; Wan et al. 2018; Engler et al. 2021; Alves et al. 2024).

Consequently, the pattern of climatic niche evolution evidenced among lineages of *Phyllotis* studied can contribute to advance understanding about the scenarios for diversification responsible of the diversity of the *P. darwini* group.

## 2 MATERIALS AND METHODS

### 2.1 Species distribution and bioclimatic data

A dataset of 232 occurrences was obtained from previous studies that examined species limits and update geographic distributions for *Phyllotis* species in the *darwini* group (e.g., Steppan & Ramírez 2015; Jayat et al. 2016, 2021; Teta et al. 2018, 2022; Ojeda et al. 2021; Storz et al. 2020, 2024; Quiroga-Carmona et al. 2025). Subsequently, an additional dataset of 6083 occurrences was gathered from the Global Biodiversity Information Facility (GBIF), by implementing the *rgbif* R package (Chamberlain et al. 2017). These datasets were curated by projecting each record over the most updated species ranges (Quiroga-Carmona et al. 2025), and verifying that records correspond to voucher specimens. Simultaneously, geographically redundant entries obtained for some species (*P. darwini, P. vaccarum*, and *P. xanthopygus*) were removed to keep a single record per each locality. Occurrences from areas where species ranges overlap were only preserved when it was possible to corroborate their taxonomic identity or were already examined in previous studies. Once these steps were completed, the occurrence datasets were merged, resulting in a total of 1129 records. Given that several records were located less than 2 km apart, a spatial thinning was applied using the *spThin* R package (Aiello-Lammens et al. 2015) to prevent potential biases caused by spatial autocorrelation and oversampling of environmental conditions (Boria et al. 2014). This procedure reduced the final dataset to 982 records.

Occurrences of each species were projected onto the geographic space using the QGIS software, version 3.28 Firenze (Quantum GIS Development Team, 2023), and circles with 50 km of radius were superimposed on these points, which were later merged and dissolved to establish an enclosing polygon for each species (Figure 1). Extensions of these polygons were defined considering the geographic location and accessibility of the potentially suitable areas for each species and the existence of physical barriers (e.g., lowland valleys or small ranges) to dispersion (Anderson & Raza 2010; Barve et al. 2011). Furthermore, in cases where species have contiguous or adjacent distributions, areas of overlap were further increased to prevent truncations during the characterization of the climatic niche and to ensure the inclusion of environmental configurations that are equivalent and/or accessible among compared species (see Saupe et al. 2018; Brown & Carnaval 2019).

The climatic niches were characterized by using the rasterized bioclimatic variables gathered from the WorldClim database (current conditions: 1970-2000), version 2.1 (Fick & Hijmans 2017), at a spatial resolution of 30 arc-second (∼1 km). The *raster* R package (Hijmans et al. 2015) was employed to stack all bioclimatic variables along with the constructed enclosing polygons. Posteriorly, climatic values registered in each spatial pixel were extracted. This climatic dataset was analyzed with the R packages *vegan* (Oksanen et al. 2020) and *caret* (Kuhn 2019) to identify variables with levels of correlations higher than |r|< 0.75, by employing a matrix of correlation based on Pearson’s correlation coefficient. Thereafter, only the following eight low-correlated variables were retained to perform subsequent analyses: annual mean diurnal range (Bio 2), temperature seasonality (Bio 4), minimum temperature of coldest month (Bio 6), mean temperature of wettest quarter (Bio 8), precipitation of wettest month (Bio 13), precipitation of driest month (Bio 14), precipitation seasonality (Bio 15), and precipitation of coldest quarter (Bio 19). Several of these variables form part of the climatic conditions that have been identified as important determinants of the geographic distribution of several species of *Phyllotis* (see Ruiz-Barlett et al. 2019; López-Berrizbeitia et al. 2025).

### 2.2 Characterizations and comparisons of climatic niches

Two multidimensional approaches were used to characterize the climatic niches of the species in the *Phyllotis darwini* group and to evaluate whether differences or similarities exist that could be considered as proxies of niche divergence, or conversely, niche conservatism, respectively. The first approach was focused on the realized (*sensu* Peterson et al. 2011) or occupied (*sensu* Brown & Carnaval 2019) climatic niche, as it involved sampling the climatic conditions present at the localities where species have been recorded. The second approach, in addition to considering the previously signaled portion of the climatic niche, also considers the fundamental or potential climatic niche (see Brown & Carnaval 2019), as climatic conditions present in the background areas (the geographic areas that extends beyond the observed distribution of species; Jezkova & Wiens 2018)—and therefore broader in climatic terms—were also characterized (Peterson et al. 2011; Brown & Carnaval 2019). This second approach assumes that the species’ current geographic distributions are in a non-equilibrium state with respect to the occupation of the accessible environmental suitable areas. Consequently, it can identify whether observed niche differences occur among the fundamental niches or are those merely resulting from differences in accessibility to environmental conditions present in the species’ geographic ranges (see Brown & Carnaval 2019). Assessing differences among species of *Phyllotis* based on both approaches is relevant because their realized climatic niches may comprise different portions within the same accessible environmental space that encompasses their fundamental climatic niches (see Soberón & Nakamura 2009). This implies that differences among the species’ realized climatic niches may actually be due to the inaccessibility of the specific portions of the environmental spaces, rather than due to differences associated with their climatic preferences or own abilities to occupy these climatic spaces (see Wielstra et al. 2012; Brown & Carnaval 2019).

To implement the first approach, the *raster* R package was employed to stack the selected bioclimatic variables with the occurrence points of each species, and to extract the values of all climatic parameters at their respective localities. This climatic dataset was analyzed using a Principal Component Analysis (PCA) performed with the *ade4* R package (Dray & Dufour 2007), following the technique described by Broennimann et al. (2012) as PCA-occ. Once PCA-occ scores are plotted in a common environmental space defined by the orthogonal projection of PCA axes, niche conservatism or divergence among *Phyllotis* species can be assessed by evaluating the degree of overlap or segregation of their realized climatic niches (Broennimann et al. 2012). Statistical significance of niche differences was assessed with a Pairwise Permutational Multivariate Analysis of Variance (PERMANOVA) performed with 1×10^4^ iterations and a Euclidean distance matrix calculated from the scores of all PCs using the *vegan* R package. Since the PCA-occ and PERMANOVA results do not provide direct measures of niche similarity and overlap, PCA-occ scores were also used to quantify these parameters by calculating the intersections among niche hypervolumes in a multidimensional space. This quantification provides a more geometrically realistic representation of the realized niche in a Hutchinsonian perspective, as it accounts for the extreme values of environmental dimensions and for the density of occurrences, which are essential aspects for accurately quantifying niche differentiation among species (Blonder et al. 2014, 2018; Mammola 2019). Dimensions of PCA-occ were reduced by selecting the most relevant principal components (PCs) using the Broken Stick model of MacArthur (1957) and the R function *evplot* (Gupta 2019). Thus, the scores of the first four PCs, which account for 87% of the explained variance (see Table S1 in Supplementary Materials), were used to calculate Gaussian convex hypervolumes representing the realized climatic niche of each species (Blonder et al. 2014, 2018). These ellipsoids were then simultaneously projected into a common climatic space to evaluate their degree of uniqueness, segregation, and/or overlap (see Carvalho & Cardoso 2020), using the *hypervolume* R package (Blonder et al. 2014, 2018). Pairwise comparisons between species’ ellipsoids were conducted using the Jaccard and Sørensen indices as metrics of niche similarity, with values ranging from 0 (fully dissimilar) to 1 (fully similar), describing the proportion of the climatic niche space shared by the compared species (Mammola 2019). Uniqueness fractions were also calculated for each comparison, corresponding to the fraction of each ellipsoid that is unique to each species and whose values (0 to 1) are interpreted in the same way.

The second approach contemplates characterizing both realized and fundamental climatic niches. Therefore, climatic values across the environmental space delimited by the species’ background areas were also sampled. Hence, the *raster* R package was used to project the species polygons onto the selected bioclimatic variables and to extract the climatic values from each pixel enclosed by them. These pixel-based characterizations, together with those completed from the occurrence points, were analyzed with the *humboldt* R package (Brown & Carnaval 2019) to evaluate whether differences between the climatic niches of 11 species composing the *darwini* group are statistically significant. With this approach the total environmental data characterized for species is reduced to two dimensions, standardized by means of a PCA defined by Broennimann et al. (2012) as PCA-env. After this procedure, the niche overlap test (hereafter NOT) and niche divergence test (hereafter NDT) were applied. NOT compares species’ niches across their entire geographic ranges by evaluating the multidimensional environmental space (E-space) available within the geographic region considered (G-space) (Brown & Carnaval 2019). This analysis uses the full set of accessible environmental areas to the species being compared, and statistically significant differences (*p*-values ≤ 0.05) detected indicate that climatic niches are non-equivalent (NNE), but do not allow discrimination of whether this is due to real differences in the species’ fundamental niches or simply due to access to non-analogous portions of the shared environmental space. In contrast, NDT focuses on environmental conditions that are accessible and shared by the species being compared (i.e., analogous accessible environment). This analysis therefore tests whether observed niche differences reflect true divergence resulting from differentiation of fundamental climatic niches, rather than differences leading on occupations of different portions of the shared environmental space. When NDT results are statistically significant, demonstrates that species occupying analogous accessible environments still maintain non-equivalent niches, providing evidence for divergence of their niches (ND), rather than differences driven solely by different access to this shared environmental space (Brown & Carnaval 2019). Finally, if neither test reveals significant differences, the species are considered to share similar climatic tolerances and to have equivalent niches (NE). During the development of theses analyses niche overlap was also quantified using Schoener’s D. This metric ranges from 0 (indicating complete divergence/disjunction of species niches) to 1 (indicating complete conservation/overlap of species niches). Resulting values were categorized into five discrete intervals of niche overlap, following Rödder & Engler (2011): no overlap (0-0.2), low overlap (0.2-0.4), moderate overlap (0.4-0.6), high overlap (0.6-0.8), and very high overlap (0.8-1.0).

A last approach was applied to evaluate patterns of climatic niche overlap among species and their climatic tolerances along each bioclimatic variable, examining the independent relevance of each environmental dimension in structuring the climatic niche of each species (see Engler et al. 2021), but without unifying all variables into a multidimensional space (i.e., in a Hutchinsonian sense). This third approach was developed by constructing profiles of predicted niche occupancy (PNO) for each species along each bioclimatic variable, following the framework described by Evans et al. (2009) and using the *phyloclim* R package (Heibl & Calenge 2013). Quantifications of species climatic tolerances were estimated by modeling the climatic niche of each species. Thus, a climatic niche model (CNM) was developed for each species of *Phyllotis* using the maximum entropy method implemented in MaxEnt, version 3.4.4. (Phillips et al. 2006, 2018), the selected bioclimatic variables, and the occurrences gathered for each species. MaxEnt CNMs were constructed using default algorithm configuration, and raw probabilities of environmental suitability were combined to the bioclimatic variable values to obtain a unit area histogram of suitability, which denotes the tolerance of the species along a given climatic dimension (see Evans et al. 2009). The performance of each CNM was evaluated using the area under the receiver operating characteristic curve (AUC) and the omission rate under the tenth percentile of training presence (OR). According to Merow et al. (2013) and Shcheglovitova & Anderson (2013), low OR values indicate minimal overfitting to the calibration dataset, whereas high AUC values suggest greater discrimination power between occurrences and background records. Subsequently, the degree of interspecific overlap along each bioclimatic variable was quantified using Schoener’s D as metric of niche overlap.

### 2.3 Evolutionary analyses of climatic niches

More recent approaches aimed at assessing climate niche evolution, either to explore conservatism or divergence, have prioritized characterizations of climate envelopes across geographic space (over those based solely on minimum, maximum, or mean values) given the inability of these metrics to capture the inherent multimodal variability of climate and its frequent lack of normality (see Evans et al. 2009; Nyári & Reedy 2013; Budic & Dormann 2015). Considering this caveat, the evolution of climatic niches among species of the *darwini* group was assessed by integrating a dated phylogeny with climatic data gathered after characterizing the climatic niche of each species and constructing the PNO profiles. This information was employed with two main purposes: to explore the potential association between phylogenetic relatedness of species and their climatic niche dissimilarities, and to reconstruct the evolutionary trajectories of occupations realized by each species within particular intervals in each climatic dimension. The dated phylogenetic tree inferred by Quiroga-Carmona et al. (2025) using a fragment of the mitochondrial cytochrome-b gene (Cytb) for the species of *Phyllotis* was pruned using the *ape* R package (Paradis & Schliep 2019), to retain in a more reliable way the phylogenetic relationships among the 11 species of the *darwini* group and the reconstructed evolutionary rates (Ryberg & Matheny 2011). The chronogram shown in Figure 1 depicts the evolutionary relationships and ages of divergence of these 11 species. Most of these relationships were already recovered through analyzing the complete mitochondrial genomes (Quiroga-Carmona et al. 2025) and also were inferred by other phylogenetic studies (e.g., Steppan et al. 2004, 2007; Ojeda et al. 2021; Jayat et al. 2016, 2021, 2022; Storz et al. 2024).

Patterns of evolution related to variations in the bioclimatic dimensions that structure the niche of each species were initially examined by fitting three evolutionary models (Brownian motion, BM; Early burst, EB; Ornstein-Uhlenbeck, OU) to their median values, considering that that this statistic is a robust representation of the probability density of a biological population whose natural lower bound is zero (Ruiz-Barlett et al. 2019). This process was done with the *geiger* R package (Pennell et al. 2014); the best model fit was selected using the corrected Akaike information criterion (AICc) and the Akaike weights (ω). Subsequently, the phylogenetic signal across bioclimatic variables was evaluated by inputting their median values and the calibrated tree in the *phylosignal* R package (Keck et al. 2016), running 1000 random subsamples and employing five different metrics (Abouheif’s Cmean, Moran’s I, Blomberg’s K and K*, as well as Pagel’s λ). Finally, phylogenetic correlograms were computed for changes across each bioclimatic variable to test the Local Indicator of Phylogenetic Association (LIPA; Anselin 1995) in each tip of the date tree, using the ‘two-sided’ alternative hypothesis. These analyses were aimed to assess whether similarities among climatic traits defining niches (i.e., the distribution of the occupancy of a bioclimatic variable by a species) tend to be greater among closely related species than among unrelated ones. The phylogenetic signal would provide a general overview of this, whereas the results of the LIPA would indicate where, across the phylogeny, the phylogenetic associations of climatic niche variations are or are not statistically significant (Keck et al. 2016).

Climatic niche evolution was also explored for each species following the methodology described in Engler et al. (2021). Reconstructions of ancestral climatic tolerances in G-space (i.e., considering the geographic context in which the niche climatic conditions exist) were developed to provide evidence of signals of conservatism or divergence of climatic niches under the assumption of Brownian motion evolution (Losos 2008; Budic & Dormann 2015). This estimation was performed with 1000 resampling iterations of the weighted mean calculated from PNO profiles and the GLS method by implementing the *phyloclim* R package. For the ancestral reconstructions of climatic niches in E-space (i.e., only considering the environmental space in which these exist), the median values of each climatic variable were calculated from the values extracted from the occurrences, a Brownian motion model of evolution was set in the *anc*.*ML* function of the *phytools* R package (Revell 2012), as this was the model that best fit the variation of all bioclimatic variables (a complete overview on model selection is provided in Table S2 in SM). Uncertainty in reconstructions was assessed with 75% of randomly selected information for each species across each bioclimatic variable, completing 200 repetitions, and results were plotted on phenograms to represent the niche evolution trajectories based on the best-fit evolutionary model selected. All analytical procedures were carried out using the R platform, version 4.1.0 (R Core Team 2021) and functions available in the R packages used.

## 3 RESULTS

### 3.1 Comparisons of climatic niches

The PCA-occ approach indicated high levels of differentiation among the realized climatic niches of species in the *Phyllotis darwini* group. None of the sister species pairs exhibited levels of niche overlap exceeding one-tenth (i.e., values higher than 0.1 in Jaccard or Sørensen indices) of the multidimensional space of their realized climatic niches. In these comparisons, values of niche uniqueness were larger than 0.8 (Table 1). This general pattern of differentiation among species’ realized climatic niches holds true among most species of *Phyllotis* with the exception of two pairwise comparisons: *P. limatus* versus *P. chilensis*, and *P. limatus* versus *P. magister* (cases 29 and 41, respectively, in Table S3). The PCA-occ provided an effective summary of climatic niches, as the first three PCs captured more than 75% of the variance contained in the bioclimatic variables selected (Table S1). PCs loadings showed that the components are influenced almost equally by the bioclimatic variables describing temperature (Bio 2-Bio 8) as well as those describing precipitation patterns (Bio 13-Bio 19). Among these, precipitation seasonality (Bio 15), mean temperature of wettest quarter (Bio 8), precipitation of driest month (Bio 14), and minimum temperature of coldest month (Bio 6), had PCs loadings ≤|0.7|. Similarly, PERMANOVA results revealed significant differences (*p*-values ≤ 0.05) in almost all pairwise comparisons of species’ climatic niches, except for two of them: *P. limatus* versus *P. chilensis* and *P. limatus* versus *P. magister* (Table S3). Finally, projections of the convex Gaussian hypervolumes also indicated low levels of overlap between the ellipsoids representing the realized climate niche of each species (Figure 2). The Jaccard and Sørensen indices did not exceed 0.140 and 0.245, respectively, and averaged less than 0.02 (calculated without considering non-significant niche comparisons; see Table S3).

**Table 1.** Overlap and uniqueness calculated from the Gaussian convex hypervolumes representing the realized climatic niches of sister species of *Phyllotis* within the *darwini* group. Additionally, in the last columns are presented the results of the PERMANOVA performed to evaluate the significance of these comparisons. Overlap and uniqueness values range from 0 to 1; where a value of 1 indicates complete overlap or complete uniqueness, respectively. PERMANOVA results were considered significant when p-values are less than 0.05. Pseudo-F value of each pairwise comparison is also provided. The complete list of pairwise comparisons completed among all species of *Phyllotis* is presented in Table S3.

| Case | Species compared | Niche overlap |  | Niche uniqueness |  | PERMANOVA |  |
| --- | --- | --- | --- | --- | --- | --- | --- |
|  |  | Jaccard | Sørensen | U of 1<sup>st</sup> Sp. | U of 2<sup>st</sup> Sp. | p-value | Pseudo-F |
| 1 | *P. bonariensis* vs. *P. camiari* | 0.000 | 0.000 | 1.00 | 1.00 | 0.0055 | 379.527 |
| 22 | *P. caprinus* vs. *P. limatus* | 0.012 | 0.023 | 0.86 | 0.99 | 0.0055 | 16.690 |
| 26 | *P. caprinus* vs. *P. vaccarum* | 0.000 | 0.000 | 1.00 | 1.00 | 0.0055 | 38.060 |
| 32 | *P. chilensis* vs. *P. posticalis* | 0.073 | 0.137 | 0.83 | 0.89 | 0.0055 | 46.482 |
| 36 | *P. darwini* vs. *P. magister* | 0.000 | 0.000 | 1.00 | 1.00 | 0.0055 | 82.917 |
| 44 | *P. limatus* vs. *P. vaccarum* | 0.014 | 0.027 | 0.96 | 0.98 | 0.0055 | 46.742 |
| 52 | *P. pehuenche* vs. *P. xanthopygus* | 0.018 | 0.035 | 0.80 | 0.98 | 0.0055 | 15.195 |

**Figure 2.**
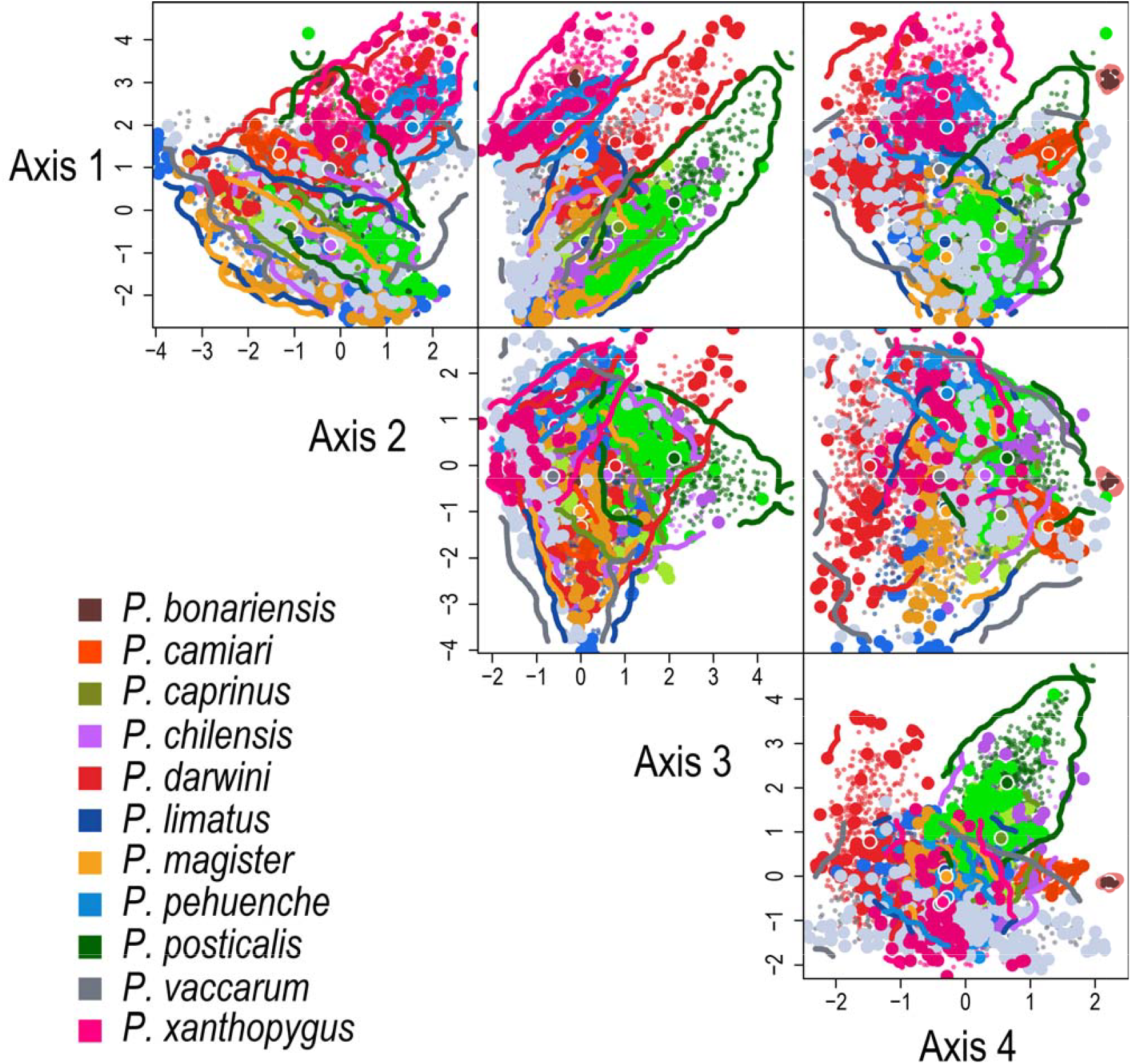
Simultaneous projection of the convex Gaussian hypervolumes in the bidimensional climatic spaces constructed using the orthogonal projections of the first four principal components obtained with the PCA-occ. These four principal components explain 87.07% of the variance (see Table S1). Each box displays an orthogonal projection of PCs, depicting the centroid (large circles with white strokes) of each convex polygon (colored contour line), the records of each species (larger dots), and the random points (smaller dots) used to calculate each species’ convex Gaussian hypervolume. Percentage of variance accounted by each PC is shown on each corresponding side. Taxon names are indicated by the name of each species of Phyllotis.

Results of climatic niche comparisons performed using NOT and NDT analyses—which are based on the PCA-env approach—were broadly consistent with the results obtained with the PCA-occ approach. Specifically, these tests also demonstrated that the fundamental climatic niches of the species within the *darwini* group exhibit high levels of differentiation. Most comparisons revealed interspecific differentiation of climatic niches, as 38 of 55 pairwise comparisons were classified as cases of niche divergence (ND), 13 were classified as niche non-equivalence (NNE), and only four were classified as niche equivalence (NE; see Tables 2 and Table S4), indicating that fundamental climatic niches have diverged between most species (Brown & Carnaval 2019). In 40 of these 55 comparisons, Schoener’s D index that did not exceed 0.2, falling in the no overlap category defined by Rödder & Engler (2011), followed by 11 cases classified as low overlap (0.2-0.4), and solely four cases classified as moderate overlap (0.4-0.6; Table S4). In all implementations of the PCA-env approach, developed to complete these analyses, the first two PCs always summarized more than 70% of the explained variance (Table S4), and the kernel density isopleths used to represent the fundamental climatic niche of each species illustrate the low levels of niche overlap observed in most comparisons (Figure 3). On the orthogonal projection of these PCs, most overlaps between species’ fundamental climatic niches occur in the lower quartiles, while those occurring in the upper quartiles are less frequent; which evidences the high differentiation between the climatic niches compared.

**Table 2.** Pairwise comparisons of the fundamental climatic niches of the sister species of *Phyllotis* within the *darwini* group. Results of these comparisons completed using the niche overlap test (NOT) and the niche divergence test (NDT) are depicted. Results of these niche comparisons were classified as niche divergence (ND), niche equivalence (NE), or as niche non-equivalence (NNE), according to guidelines described by Brown & Carnaval (2019) for the jointly interpretation of p-values (see section 2.2 in Material and Methods). Niche similarity is quantified based on Schoener’s D index of niche overlap, and p-values of statistical significance are provided. The percentage of explained variance (% EV) accounted by the two PCs used to perform each pairwise comparison is also shown. The complete list of pairwise comparisons of species’ fundamental climatic niches is presented in Table S4.

| Case | Species compared | NOT (Full Extension) |  |  |  | NDT (Shared Extension) |  |  |  | % EV |  |
| --- | --- | --- | --- | --- | --- | --- | --- | --- | --- | --- | --- |
|  |  | Niche Similarity | *p*-value | Background 2→1 | Background 1→2 | Niche Similarity | *p*-value | Background 2→1 | Background 1→2 | PC 1 | PC 2 |
| 1 | *P. bonariensis* vs. *P. camiari*<sup>ND</sup> | 0.0000 | 0.0043 | 0.0099 | 0.0099 | 0.0000 | 0.0105 | 0.0099 | 0.0099 | 52.13 | 24.87 |
| 22 | *P. caprinus* vs. *P. limatus*<sup>NNE</sup> | 0.2130 | 0.0453 | 0.0287 | 0.0292 | 0.2160 | 0.0573 | 0.0213 | 0.0233 | 50.82 | 23.78 |
| 26 | *P. caprinus* vs. *P. vaccarum*<sup>ND</sup> | 0.0740 | 0.0099 | 0.0594 | 0.0297 | 0.0710 | 0.0099 | 0.0257 | 0.0614 | 49.01 | 29.60 |
| 32 | *P. chilensis* vs. *P. posticalis*<sup>NNE</sup> | 0.4110 | 0.0289 | 0.0491 | 0.0491 | 0.4110 | 0.0871 | 0.0491 | 0.0495 | 64.26 | 15.42 |
| 36 | *P. darwini* vs. *P. magister*<sup>ND</sup> | 0.0300 | 0.0089 | 0.0478 | 0.0472 | 0.0310 | 0.0099 | 0.0432 | 0.0212 | 47.70 | 26.81 |
| 44 | *P. limatus* vs. *P. vaccarum*<sup>NNE</sup> | 0.3490 | 0.0241 | 0.0589 | 0.0589 | 0.3330 | 0.0576 | 0.0493 | 0.0411 | 49.98 | 26.14 |
| 52 | *P. pehuenche* vs. *P. xanthopygus*<sup>NNE</sup> | 0.3770 | 0.0481 | 0.2673 | 0.0663 | 0.3770 | 0.0921 | 0.0911 | 0.0861 | 46.84 | 23.79 |

**Figure 3.**
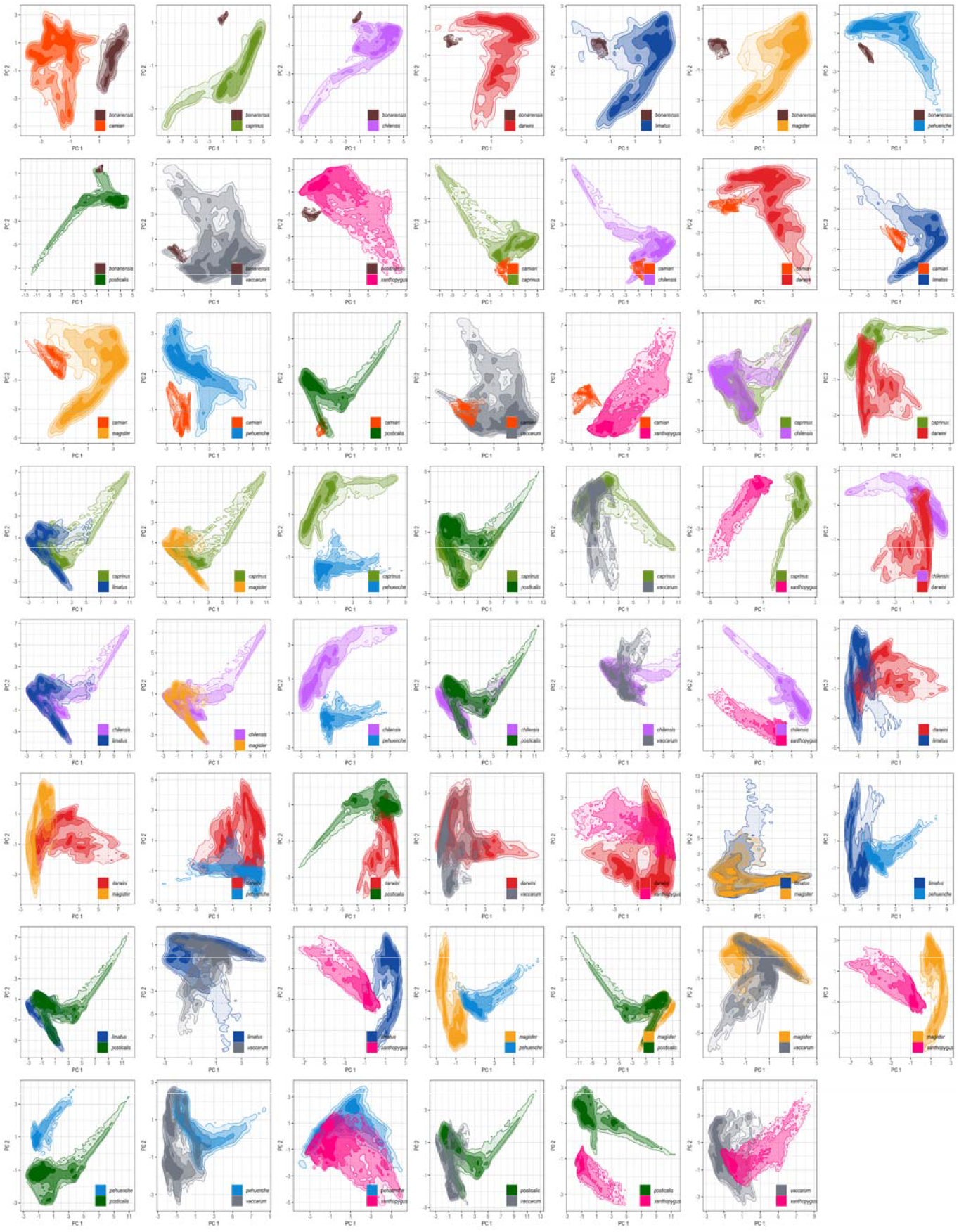
Projection of the kernel density isopleths calculated for each species of *Phyllotis* after conducting paired comparisons of their potential climatic niches with the humboldt R package. Each shaded colored area represents the total environmental space occupied by each species, with darker areas indicating higher occurrence density. The non-transparent area corresponds to the upper 50% of the density distribution, while the remaining contour lines delimit 100% of the occupied climatic space. Taxon names are indicated by the epithet of each species of *Phyllotis*. The percentage of explained variance accounted by the PC projected is provided in Table S4.

Finally, quantifications of niche overlaps based on species tolerances to each bioclimatic variable revealed that the levels of interspecific overlap are higher and more variable when they are evaluated individually along each climatic dimension than when assessed using a multidimensional approach. In these evaluations, values of Schoener’s D ranged from 0.010 to 0.988 (Tables 3 and S5), exhibiting a high variability in terms of interspecific overlap along climatic dimensions. On average, the highest level of overlap was observed along annual mean diurnal range (Bio 2), whereas occupation tended to be more scattered along temperature seasonality (Bio 4; Table S5). The PNO profiles show that levels of overlap along the other climatic dimensions are similar (Figure 4), but separation of species peaks representing the highest levels of suitability for each species along climatic dimensions are visually more evident for temperature seasonality (Bio 4), mean temperature of wettest quarter (Bio 8), and precipitation seasonality (Bio 15). Average AUC and OR values for the MaxEnt CNMs developed were 0.981 and 0.036, respectively, indicating their strong performance according to the minimal overfitting and the discrimination power (Figure S2). Furthermore, the spatial projections of these models provided a proper representation of species’ potential distributions and estimate with fidelity their observed geographic distributions (Figures 1 and S2).

**Table 3.** Climatic niche overlaps among sister species of *Phyllotis* within the *darwini* group across the uncorrelated bioclimatic variables. Quantifications of niche overlap are based on Schoener’s D metric and were completed based on the profiles of predicted niche occupancy (PNO). Values Schoener’s D of niche overlap are only show for pairwise comparisons corresponding to pairs of sister species. The complete list of comparisons is shown in Table S5.

| Case | Species compared | Bio 02 | Bio 04 | Bio 06 | Bio 08 | Bio 13 | Bio 14 | Bio 15 | Bio 19 |
| --- | --- | --- | --- | --- | --- | --- | --- | --- | --- |
| 1 | *P. bonariensis* vs. *P. camiari* | 0.873 | 0.819 | 0.898 | 0.751 | 0.914 | 0.571 | 0.659 | 0.597 |
| 22 | *P. caprinus* vs. *P. limatus* | 0.907 | 0.853 | 0.834 | 0.905 | 0.933 | 0.940 | 0.866 | 0.934 |
| 26 | *P. caprinus* vs. *P. vaccarum* | 0.954 | 0.801 | 0.899 | 0.872 | 0.833 | 0.881 | 0.786 | 0.858 |
| 32 | *P. chilensis* vs. *P. posticalis* | 0.928 | 0.726 | 0.929 | 0.895 | 0.785 | 0.831 | 0.839 | 0.852 |
| 36 | *P. darwini* vs. *P. magister* | 0.640 | 0.607 | 0.848 | 0.883 | 0.864 | 0.838 | 0.730 | 0.427 |
| 44 | *P. limatus* vs. *P. vaccarum* | 0.967 | 0.529 | 0.836 | 0.902 | 0.881 | 0.932 | 0.884 | 0.898 |
| 52 | *P. pehuenche* vs. *P. xanthopygus* | 0.898 | 0.948 | 0.967 | 0.945 | 0.944 | 0.965 | 0.885 | 0.934 |
|  | Average of niche overlap | 0.881 | 0.755 | 0.887 | 0.879 | 0.879 | 0.851 | 0.807 | 0.786 |

**Figure 4.**
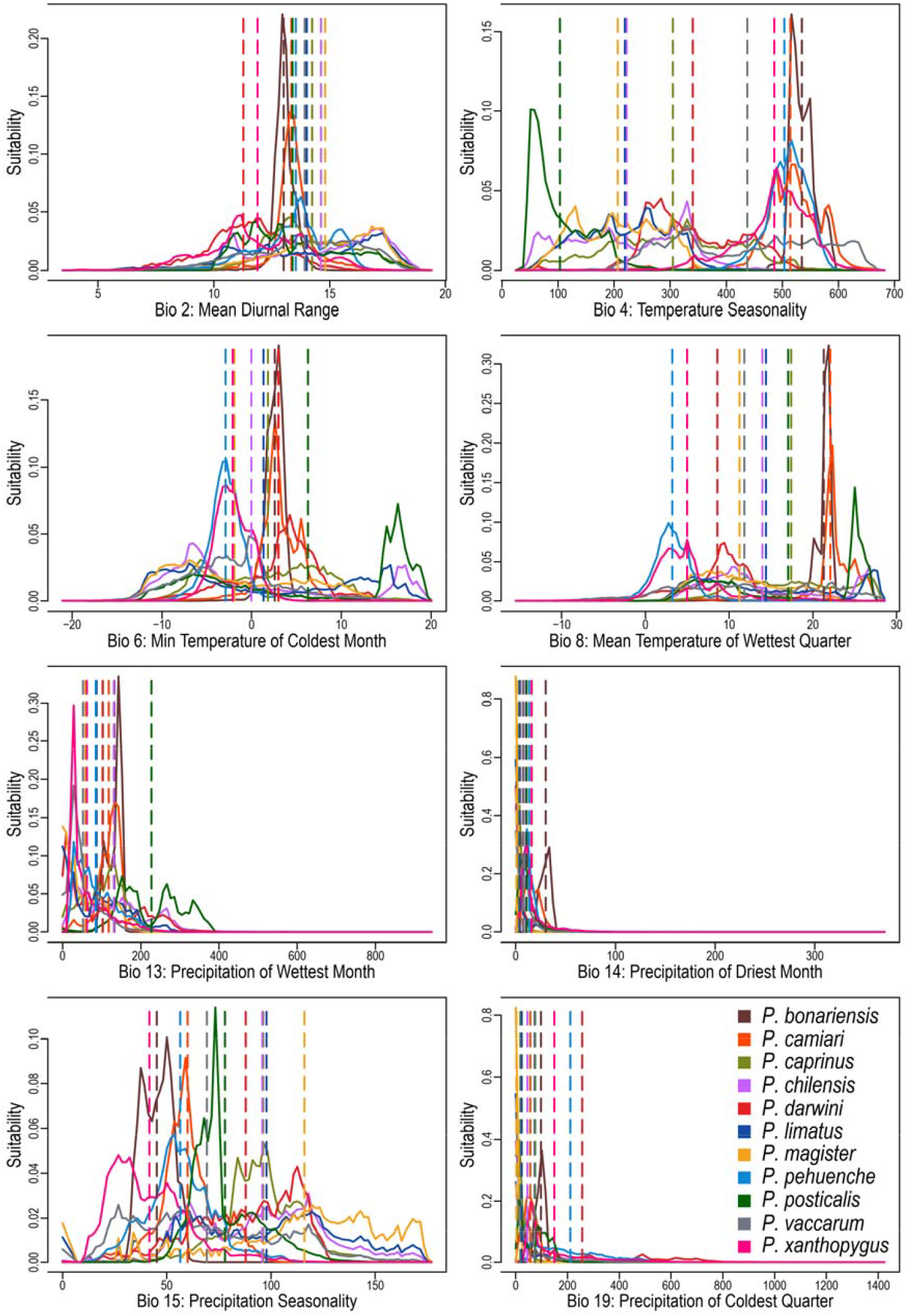
Predicted niche occupancy (PNO) profiles estimated for species of *Phyllotis* within the *darwini* group. Each panel illustrates the total suitability (y-axis) estimated for each species throughout the geographic variability of the bioclimatic variable (x-axis) evaluated. The dashed lines represent the weighted means calculated for each species on each variable. The overlapping of peaks in the profiles suggests similar climatic tolerances between the species, while the width of the profile indicates the specificity of the climatic tolerance.

### 3.2 Evolution of climatic niches and reconstructions of climatic tolerances

Among the evolutionary models considered, the Brownian motion (BM) was identified as the one that best-fit for niche variations characterized among all species, in relation to the climatic features characterized and the occupancy of the eight considered bioclimatic variables (Table S2). These variations did not show statistically significant phylogenetic signals, regardless of the metric considered or the variable examined (Table S6). None of the simulations developed to compare the performance of phylogenetic signal metrics yielded consistent evidence of statistical association among variations in climatic dimensions and the phylogenetic relatedness of species (Figure S3), even when higher percentages of Brownian motion were simulated (Figure S4). Evolutionary assessments based on correlograms evidenced negative correlations at small phylogenetic distances in four of the eight bioclimatic variables evaluated, and only one of these conditions exhibited a positive correlation (Figure S5). These results align with the patterns revealed by the LIPA, which only depicted a statically significant negative association in annual mean diurnal range (Bio 2) for *Phyllotis chilensis* and in precipitation of coldest quarter for *P. darwini* (Figure S6).

Phylogeny-based reconstructions of ancestral climatic tolerances (Figure 5) and ancestral climatic niches (Figure 6) consistently showed a high degree of differentiation in the average occupancy of bioclimatic variables among members of the *darwini* group. Degrees of differentiation vary among specific lineages, bioclimatic variables, and the dimensional spaces considered (G and E), but overall, niche divergences between sister species are clearly indicated by the fact that in all cases the lineages resulting from these cladogenetic events end up occupying separate positions along the same climatic dimension (Figures 5 and 6). In these reconstructions, the highest level of climatic niche divergence was observed between *P. darwini* and *P. magister*. In general, all evidenced niche differentiations do not show a clustering pattern in which, for example, specific sets of species consistently group together and occupy specific sections within the climatic dimensions. On the contrary, what is repeatedly observed is that sister species always occupy separate portions along the same climatic dimension and cases of niche overlapping mainly occur between non-sister lineages.

**Figure 5.**
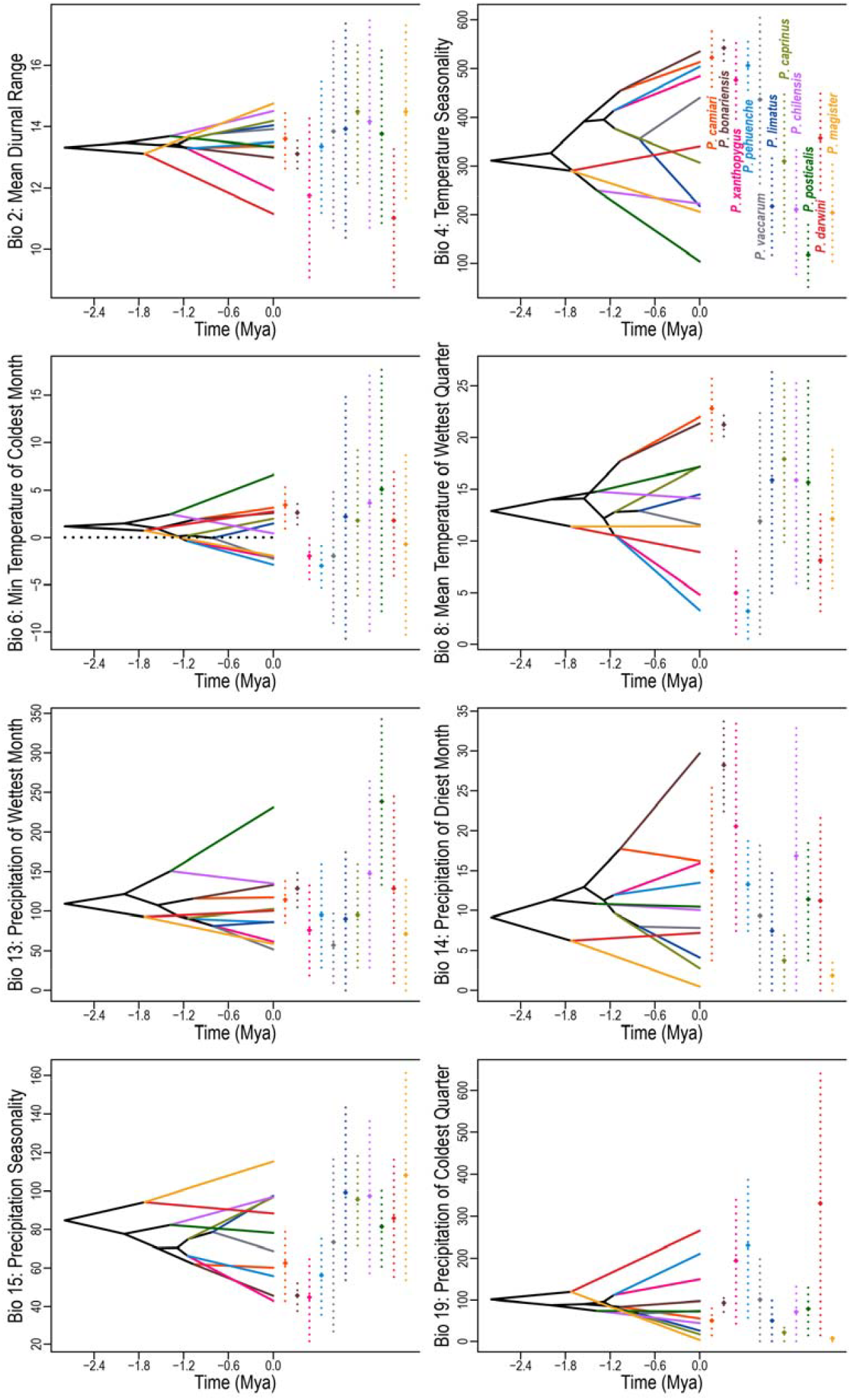
Reconstruction of ancestral climatic tolerances in G-space among species of *Phyllotis* within the *darwini* group. The phenogram is projected into niche parameter space (y-axis) through 1000 random replications of the PNO profiles and the integration of the phylogenetic tree. Internal nodes denote the weighted mean of climatic tolerances as inferred for the most recent common ancestor of the related extant taxa. Solid black lines connect ancestors with their descendants (colored lines). Vertical dashed lines indicate the 85% central density of climate tolerance for each species and diamonds indicate the weighted means.

**Figure 6.**
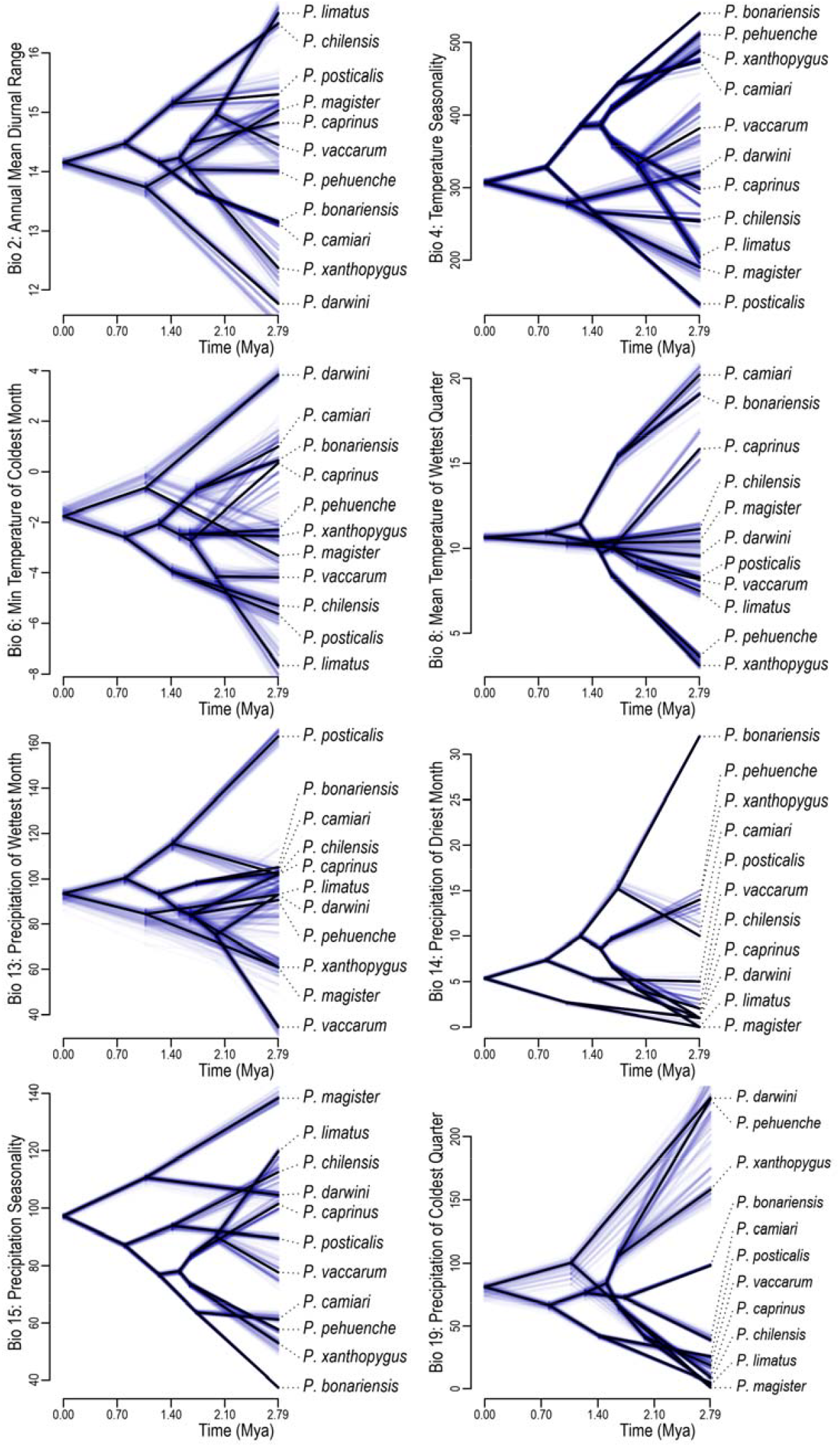
Reconstruction of the ancestral niches in E-space among species of *Phyllotis* within the *darwini* group. Ancestral niches were estimated using the trait evolution model that best fits the data (i.e., Brownian motion), and by assessing uncertainty with 200 random subsets of climatic values, constructed by incorporating 75% of the occurrence records. These estimations (phenograms shaded in blue) were projected onto the niche parameter space (y-axis). Subsequently, the full-data model (black phenogram) developed to reconstruct the ancestral niches and to represent the estimated median values at the node locations was also projected onto this space.

## 4 DISCUSSION

Analysis of climatic niche differentiations that occurred during the radiation of species in the *Phyllotis darwini* species group revealed extensive divergence in realized climatic niches (Table S3), and full divergence in fundamental climatic niches between all pairs of sister species (Table 2). In addition, the characterized patterns of climatic niche overlap indicated that niche similarities tend to be greater when quantified individually along each climatic dimension than when quantified using an integrative multidimensional approach. At first glance, these results suggest that climatic niche differences are driven primarily by the way in which these mice jointly and multidimensionally occupy climatic space, rather than by the particular use that each species makes of individual bioclimatic variables. As reflected by the absence of phylogenetic signal in climatic niche variations, the relatedness of these species was not predictive of the similarities among their climatic niches. In contrast, niche differences were even larger or smaller (see Tables S3 and S4) than would be expected, considering, for example, a constant rate of niche evolution associated with the magnitude of species divergences (e.g., as expected under a Brownian motion model, see Losos 2008, Revell et al. 2008). Overall, these results suggest that niche differentiation among species in the *darwini* group may stem largely from a history of divergent selection (see Revell et al. 2008; Münkemüller et al. 2015), likely driven by the contrasting ecoclimatic conditions prevailing in the regions where ancestral populations underwent their initial divergences in spatial isolation (Rundell & Price 2009; Sobel et al. 2010; Budic & Dorman 2015; Brown & Carnaval 2019).

### 4.1 Differentiation of climatic niches and absence of niche conservatism

Statistical analyses of the climatic niches of species in the *Phyllotis darwini* group revealed no evolutionary trend of niche conservatism. Pairwise comparisons of climatic niches consistently demonstrated that sister species occupy different portions of the available climatic space (Figures 2 and 3). The greatest levels of climatic niche overlap were observed in the comparisons between *P. limatus* and *P. chilensis* and between *P. limatus* and *P. magister*, but these overlaps between these pairs of non-sister species were still low to be considered as niche conservatism cases (Table S4). Contrary to the comparisons of realized climatic niche performed with PCA-occ, PERMANOVA, and projection of Gaussian hypervolumes, some of those done with NOT and NDT were not fully conclusive about the divergence of the fundamental climate niches of some pairs of sister species, though they did recognize the non-equivalence of their available climatic envelopes (Table S4). According to Brown & Carnaval (2019), the non-equivalence between niches cannot be taken as a direct indication of niche divergence, because access to a shared environmental space—and not the species’ climatic preferences themselves— can vary among species. In cases where the equivalence statistic is significant (*p*-values ≤ 0.05)— regardless of the significance of the background statistic—it is possible to reject the null hypothesis of niche equivalence and support the idea that species’ climatic niches are different. Thus, results indicating non-equivalence only demonstrate that the climatic spaces occupied by the species being compared are statistically different, but it cannot be concluded that these have diverged in their fundamental niches (see Brown & Carnaval 2019).

Construction of the PNO profiles showed that levels of species niche overlap are higher and more variable when quantified independently for each climatic dimension than when assessed using a multidimensional approach. The similarities and differences observed among the climatic niches of *Phyllotis* species illustrate that niche singularities are determined primarily by the joint, multivariate way in which these species occupy climatic dimensions, rather than by their separate occupation of each dimension. For instance, several species pairs (e.g., cases 1–19 in Tables S4 and S5) show Schoener’s D values close to 1 when overlap is quantified variable by variable, whereas values approach zero when the same comparisons are performed in multidimensional space. These evaluations inform our understanding of how niche occupation is structured and how niche differences arose, as they provide a more detailed depiction of how each species occupies each climatic dimension as well as the entire climatic space (Figure 4). PNO profiles also revealed that patterns of overlap lack consistent positioning across climatic dimensions; instead, occupancy peaks (and their proximity to those of other species) shift in position from one dimension to another. These observations align with and reinforce conceptual views regarding the inherently singular structure of species environmental niches (see Holt 2009; Peterson et al. 2011).

Patterns of partitioning and segregation observed across bioclimatic dimensions provide information on how climate shapes the spatial disjunction of species of the *darwini* group along the ecoclimatic gradient of southern South America. Ubiquitously, climatic specializations strongly constrain regions where species can occur and how they respond to environmental changes, thus playing a central role in shaping distributional patterns (Bonetti & Wiens 2014). These constraints derive from the physiological tolerances of individuals, which can determine the breadth, position, and frequency with which species occupy particular dimensions and configurations of climate (Kearney & Porter 2009; Bozinovic et al. 2011); especially in those species inhabiting environments characterized by extreme aridity and elevation, such as some *Phyllotis* mice that occur in highland ecosystems (see Storz & Scott 2024). In cases where species exhibited some conservatism in their climatic niches, spatial overlap would be modulated by the partitioning of Eltonian niche dimensions, such as trophic level or feeding strategy, or through displacement along other non-climatic aspects of their Grinnellian niches (Soberón 2007; Czekanski-Moir & Rundell 2019; Diniz-Filho 2023). Consequently, in regions where several of *Phyllotis* species could co-occur (e.g., near the tripartite border between Argentina, Bolivia, and Chile, where the potential distributions of *P. chilensis, P. magister, P. limatus*, and *P. vaccarum* may overlap; see Figure 1), it is plausible that syntopy is facilitated by divergence in morphological and/or behavioral traits (see Brown et al. 2002; Quintero & Landis 2020). Although recent studies have begun to elucidate how physiological tolerances and dietary habits could constrain the geographic ranges of some *Phyllotis* species (Quezada et al. 2024; Bautista et al. 2026; Liphardt et al., in press), substantial gaps remain in our understanding of the ecological and behavioral mechanisms underlying niche partitioning among Andean sigmodontine rodents.

### 4.2 Evolution of climatic niches and scenarios of diversification

Under a neutral scenario of climatic niche evolution (Coelho et al. 2019)—as represented by a Brownian motion model—it would be expected that niche differentiation among species would exhibit a phylogenetic hierarchy and the rate of niche differentiation would be roughly constant across all lineages (Losos 2008; Budic & Dormann 2015; Quintero et al. 2022). Contrary to this expectation, the phylogenetic correlograms reveal no general pattern of association between phylogenetic distances and niche differences, though four climatic dimensions exhibit statistically significant correlations, but these arose at relatively short phylogenetic distances (i.e., between some pairs of sister species; see Figures S5 and S6). According to the reconstructions of ancestral trajectories of climatic tolerances and climatic niches of species within the *darwini* group (Figures 5 and 6), sister lineages tend to occupy separate portions of the available climatic niche space. Furthermore, many of the interspecific comparisons revealed levels of niche differentiation that are higher or lower than would be expected based on the phylogenetic distances of species and a constant rate of niche differentiation (see Losos 2008; Revell et al. 2008). These results suggest that the evolutionary trajectories of climatic niches have been highly independent among the studied species (Boucher et al. 2014; Münkemüller et al. 2015), suggesting that the differentiation of climatic niches of these mice has been shaped by divergent selective pressures, such as those imposed by environmental dissimilarities among regions and intense climatic dynamics (Wiens 2004; Wang et al. 2017; Rolland et al. 2018).

Determining whether the characteristics defining the ecological niches of related species have remained conserved or have differentiated over evolutionary time has been valuable for elucidating the biogeographic context in which their diversification occurred (Kozak & Wiens 2010). This approach has provided insights into the nature of isolating barriers and the mechanisms underlying lineage divergence in several biological groups (e.g., Sheu et al. 2020; Engler et al. 2021; Alves et al. 2024). In the case of the species in the *darwini* group, modes of speciation involving sympatry can be discarded because none of the sister species have largely overlapping distributional ranges (see Turelli et al. 2001; Rundle & Nosil 2005; Sobel et al. 2010), and it is unlikely that the disjunctions of their current distributional areas occurred secondarily after sympatric speciation (see Czekanski-Moir & Rundell 2019). Geographic ranges of most of sister species of *Phyllotis* are contiguous or partially overlapping along the latitudinal ecoclimatic gradient of southern South America (Figure 1), which could be suggestive of allopatric or parapatric modes of speciation (Hua & Wiens 2013, but see Sobel et al. 2010). It is likely that the divergences of ancestral lineages were induced by their spatial separation, partly because their physiological tolerances were narrower during these earliest stages (Figures 5 and 6). Subsequent changes in local ecoclimatic conditions and habitat singularities would have contributed to accentuating the differences that emerged independently among diverging population lineages as they evolved isolated in these regions with dissimilar ecoclimatic conditions (see Rundle & Nosil 2005; Rundell & Price 2009; Crisp et al. 2011).

By adopting the perspectives of Alhajeri et al. (2016) and Maestri et al. (2017), which establish that the diversification of sigmodontine rodents occurred primarily via vicariant speciation, climatic niche conservatism would be expected to be a pervasive pattern among extant species that comprise this megadiverse group (see Wiens 2004; Rundell & Price 2009; Hua & Wiens 2013). However, the general patterns of climatic niche differentiation observed among species of the *darwini* group is consistent with the scenario described by Reis et al. (2018), as reconstructions of the evolutionary trajectories of ancestral lineages show that their climatic niches were comparatively less differentiated than the levels observed among extant species. These reconstructions also indicate that climatic niche diversification occurred progressively, ultimately producing complete differentiation between the niches of extant species. The uncovered patterns of climatic niche evolution and the isolated pattern of geographic distribution of sister species of *Phyllotis* in the *darwini* group point first to a diversification scenario in which speciation was initially driven by the geographic separation of populations. Subsequently, ecoclimatic differences among regions occupied by these populations may have promoted the differentiation of their climatic niches (see Rundell & Price 2009; Sobel et al. 2010; Crisp et al. 2011). Although niche divergence may not have been the primary mechanism triggering speciation events within this speciose clade of sigmodontine mice, it accompanied and likely reinforced the evolutionary divergence of the emerging species by limiting migration between regions with different ecoclimatic configurations (see Doebeli & Dieckmann 2004). In this context, the emergence of ecological differences during speciation has traditionally been considered a consequence of divergent natural selection exerted by environmental differences between inhabited regions on populations (Rundle & Nosil 2005; Schluter 2009), which in turn is consistent with the absence of phylogenetic association observed among the climatic niche variations of these species (see Barraclough 2019).

## 5 CONCLUSIONS

This study provides the first assessment of climatic niche differentiation and evolution in leaf-eared mice in the *Phyllotis darwini* species group. Findings outlined in this study suggest that the diversification of this group initially occurred in a context of geographic isolation and retention of ancestral climatic niches in the ancestors of extant species. Subsequent environmental changes associated with the Quaternary glaciations may have promoted niche differentiation and divergence events giving rise to current species. This study underscores the importance of an ecological perspective associated with the evolution of niches in the radiation of an especially diverse group of sigmodontine rodents, and is consistent with a role for historical climatic dynamics in promoting niche differentiation and speciation. Inferences drawn in this study are based solely on the characterization of climatic niches derived from observed species distributions. Thus, a more complete assessment of species’ climatic niches should integrate these climatic characterizations with experimental measurements of physiological tolerances directly conducted on individuals from different populations of these species. Likewise, assessments that fully consider the taxonomic composition of specific groups within the subfamily Sigmodontinae are needed to provide the necessary resolution to clearly reveal the evolutionary patterns that emerged during the diversifications of more speciose groups.

## AUTHOR CONTRIBUTIONS

**Marcial Quiroga-Carmona:** conceived and designed protocols to obtain and curate data, performed the analyses, interpreted the results, prepared figures, tables and supplementary materials, authored and reviewed drafts of the article, and approved the final version. **José H. Urquizo:** validation, visualization, writing, review and editing. **Naim M. Bautista:** validation, visualization, writing, review and editing. **Guillermo D’Elía:** validation, visualization, writing, review and editing. **Jay F. Storz:** validation, visualization, writing, review and editing.

All authors have read and approved the final version of the manuscript submitted for publication and agree to be accountable for all aspects of the work in ensuring that questions related to the accuracy or integrity of any part of the work is appropriately investigated and resolved. All persons designated as authors qualify for authorship, and all those who qualify for authorship are listed.

## ACKNOWLEDGEMENTS

To all developers of the R packages used in this study, whose work has made and will continue to make studies like this possible. No permits were required for the development of this research.

