## SUPPLEMENTARY MATERIALS for "CLIMATIC NICHE DIFFERENTIATION ACCOMPANIED THE RADIATION OF LEAF-EARED MICE IN THE *PHYLLOTIS DARWINI* SPECIES GROUP (SIGMODONTINAE, CRICETIDAE)"

***Results of Principal Component Analyses (PCA-occ)***

Results of the principal component analysis (PCA-occ) completed with the climatic characterization performed based on the climatic conditions sampled at presence points of the species within the *Phyllotis* *darwini* species group.

**Table S1.** Results of the PCA-occ performed based on the climatic conditions sampled at the presence records collected for the species of *Phyllotis* studied here. Contributions of the bioclimatic variables to the first four principal components are provided, together with eigenvalues and cumulative percentages of explained variance (% EV).

| Bioclimatic variables | PC 1 | PC 2 | PC 3 | PC 4 |
| --- | --- | --- | --- | --- |
| Bio 2: Annual mean diurnal range | -0.677 | 0.375 | -0.086 | 0.536 |
| Bio 4: Temperature seasonality | 0.666 | 0.133 | -0.588 | 0.189 |
| Bio 6: Minimum temperature of coldest month | 0.485 | -0.756 | 0.328 | -0.127 |
| Bio 8: Mean temperature of wettest quarter | 0.067 | -0.800 | 0.201 | 0.507 |
| Bio 13: Precipitation of wettest month | 0.101 | 0.516 | 0.804 | 0.198 |
| Bio 14: precipitation of driest month | 0.793 | 0.296 | 0.069 | 0.395 |
| Bio 15: Precipitation seasonality | -0.810 | -0.002 | 0.138 | -0.190 |
| Bio 19: precipitation of coldest quarter | 0.694 | 0.428 | 0.196 | -0.319 |
| Eigenvalues | 2.920 | 1.907 | 1.209 | 0.929 |
| Cumulative percentages of explained variance | 36.499 | 60.338 | 75.452 | 87.068 |

Principal components were considered relevant when percentages of explained variance, calculated after standardizing the data, surpassed the values of variance predicted by the Broken Stick model; which indicated that the variance they represent is greater than expected under a null model of random allocation of variance. The result of this selection is shown in Figure S1, presented below.

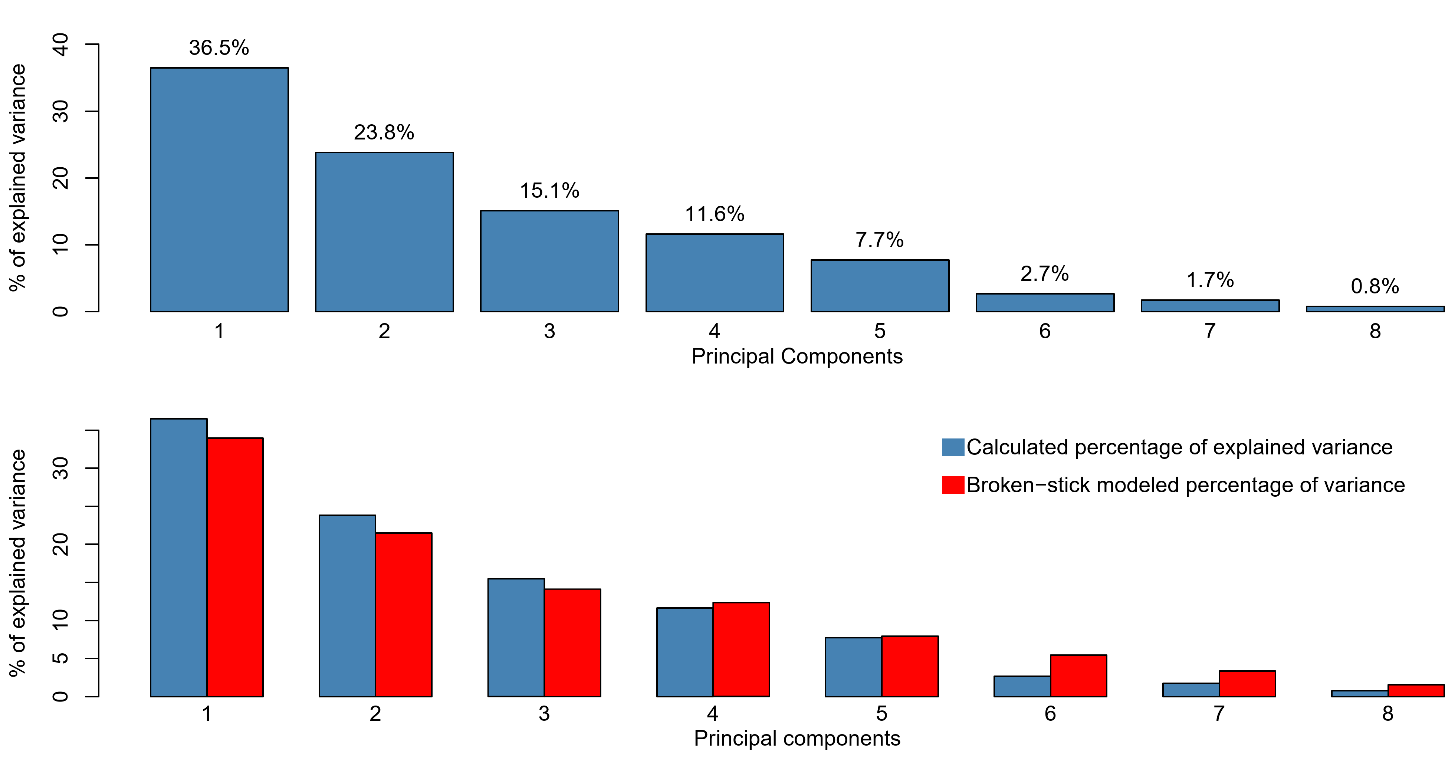

**Figure S1.** Implementation of the Broken Stick model to select the principal components considered relevant for statistical analyses of the realized climatic niches. Blue bars represent the observed values of variance and red bars represented the modeled values of explained variance according to a null model.

***Fitting of Evolutionary Models***

Median values calculated for each species within the *Phyllotis* *darwini* species group across each of the eight bioclimatic variables were fitted to three evolutionary models: Brownian motion, Early burst, and Ornstein-Uhlenbeck. The R package geiger (Harmon et al. 2008) was implemented to integrated bioclimatic and phylogenetic data, and values of Napierian logarithm (-ln *L*), corrected Akaike information criterion (AICc), and Akaike weights (ω) were retained for each procedure (Table S2). Thus, the best model fitting was selected based on the corrected Akaike information criterion (AICc) and the Akaike weights (ω).

**Table S2.** Fitting of the alternative evolution models for the eight selected bioclimatic variables. Values for Napierian logarithm (-ln *L*), corrected Akaike information criterion (AICc), and Akaike weights (ω) obtained for each variable are shown. Models with best fit (i.e., those that were selected and used in evolutionary analyses and reconstructions) are those indicated in bold.

| Bioclimatic variable | Evolution Model | -ln *L* | AICc | ω |
| --- | --- | --- | --- | --- |
| Bio 2  Annual mean diurnal range | **Brownian motion** | **-20.8261** | **47.1521** | **0.7046** |
|  | Ornstein-Uhlenbeck | -20.1386 | 49.7058 | 0.1965 |
|  | Early burst | -20.8261 | 51.0807 | 0.0988 |
| Bio 4  Temperature seasonality | **Brownian motion** | **-69.4657** | **144.4315** | **0.7671** |
|  | Ornstein-Uhlenbeck | -69.3134 | 148.0555 | 0.1253 |
|  | Early burst | -69.4657 | 148.3600 | 0.1076 |
| Bio 6  Minimum temperature of coldest month | **Brownian motion** | **-28.8127** | **63.1254** | **0.7506** |
|  | Ornstein-Uhlenbeck | -28.4988 | 66.4262 | 0.1441 |
|  | Early burst | -28.8127 | 67.0539 | 0.1053 |
| Bio 8  Mean temperature of wettest quarter | **Brownian motion** | **-34.5628** | **74.6256** | **0.7209** |
|  | Ornstein-Uhlenbeck | -33.9970 | 77.4226 | 0.1780 |
|  | Early burst | -34.5628 | 78.5542 | 0.1011 |
| Bio 13  Precipitation of wettest month | **Brownian motion** | **-54.4743** | **114.4487** | **0.6673** |
|  | Ornstein-Uhlenbeck | -53.5364 | 116.5013 | 0.2391 |
|  | Early burst | -54.4743 | 118.3773 | 0.0936 |
| Bio 14  Precipitation of driest month | **Brownian motion** | **-40.7959** | **87.0917** | **0.7147** |
|  | Ornstein-Uhlenbeck | -40.1826 | 89.7937 | 0.1851 |
|  | Early burst | -40.7959 | 91.0203 | 0.1002 |
| Bio 15  Precipitation seasonality | **Brownian motion** | **-52.9004** | **111.3008** | **0.7706** |
|  | Ornstein-Uhlenbeck | -52.7850 | 114.9985 | 0.1213 |
|  | Early burst | -52.9004 | 115.2294 | 0.1081 |
| Bio 19  Precipitation of coldest Quarter | **Brownian motion** | **-65.4388** | **136.3776** | **0.6803** |
|  | Ornstein-Uhlenbeck | -64.5840 | 138.5966 | 0.2243 |
|  | Early burst | -65.4388 | 140.3062 | 0.0954 |

***Results of pairwise comparisons of the realized climatic niches of* Phyllotis *species within the* darwini *species group***

**Table S3**. Overlap and uniqueness indices calculated for the 55 cases corresponding to all pairwise comparisons completed among 11 *Phyllotis* species within the *darwini* species group. Additionally, in the last columns are presented the results of the PERMANOVA performed to evaluate the significance of these comparisons. Overlap and uniqueness values range from 0 to 1; where a value of 1 indicates complete overlap or complete uniqueness, respectively. PERMANOVA results were considered significant when *p*-values are less than 0.05. Pseudo-F values of each comparison are also provided. Comparisons completed between sister species are those represented in bold.

| Case | Species compared | Niche overlap | | Niche uniqueness | | PERMANOVA | |
| --- | --- | --- | --- | --- | --- | --- | --- |
|  |  | Jaccard | Sørensen | U of 1^st^ sp. | U of 2^st^ sp. | *p*-value | Pseudo-F |
| **1** | ***P. bonariensis vs. P. camiari*** | **0.000** | **0.000** | **1.00** | **1.00** | **0.0055** | **379.527** |
| 2 | *P. bonariensis vs. P. caprinus* | 0.000 | 0.000 | 1.00 | 1.00 | 0.0055 | 412.435 |
| 3 | *P. bonariensis vs. P. chilensis* | 0.000 | 0.000 | 1.00 | 1.00 | 0.0055 | 319.926 |
| 4 | *P. bonariensis vs. P. darwini* | 0.000 | 0.000 | 1.00 | 1.00 | 0.0055 | 132.021 |
| 5 | *P. bonariensis vs. P. limatus* | 0.000 | 0.000 | 1.00 | 1.00 | 0.0055 | 185.632 |
| 6 | *P. bonariensis vs. P. magister* | 0.000 | 0.000 | 1.00 | 1.00 | 0.0055 | 358.058 |
| 7 | *P. bonariensis vs. P. pehuenche* | 0.000 | 0.000 | 1.00 | 1.00 | 0.0055 | 316.406 |
| 8 | *P. bonariensis vs. P. posticalis* | 0.000 | 0.000 | 1.00 | 1.00 | 0.0055 | 408.077 |
| 9 | *P. bonariensis vs. P. vaccarum* | 0.000 | 0.000 | 1.00 | 1.00 | 0.0055 | 119.236 |
| 10 | *P. bonariensis vs. P. xanthopygus* | 0.000 | 0.000 | 1.00 | 1.00 | 0.0055 | 127.294 |
| 11 | *P. camiari vs. P. caprinus* | 0.000 | 0.000 | 1.00 | 1.00 | 0.0055 | 49.948 |
| 12 | *P. camiari vs. P. chilensis* | 0.000 | 0.000 | 1.00 | 1.00 | 0.0055 | 65.950 |
| 13 | *P. camiari vs. P. darwini* | 0.000 | 0.000 | 1.00 | 1.00 | 0.0055 | 31.537 |
| 14 | *P. camiari vs. P. limatus* | 0.000 | 0.000 | 1.00 | 1.00 | 0.0055 | 39.893 |
| 15 | *P. camiari vs. P. magister* | 0.000 | 0.000 | 1.00 | 1.00 | 0.0055 | 78.406 |
| 16 | *P. camiari vs. P. pehuenche* | 0.000 | 0.000 | 1.00 | 1.00 | 0.0055 | 121.469 |
| 17 | *P. camiari vs. P. posticalis* | 0.000 | 0.000 | 1.00 | 1.00 | 0.0055 | 106.608 |
| 18 | *P. camiari vs. P. vaccarum* | 0.000 | 0.000 | 1.00 | 1.00 | 0.0055 | 22.756 |
| 19 | *P. camiari vs. P. xanthopygus* | 0.000 | 0.000 | 1.00 | 1.00 | 0.0055 | 57.256 |
| 20 | *P. caprinus vs. P. chilensis* | 0.087 | 0.159 | 0.27 | 0.91 | 0.0055 | 25.738 |
| 21 | *P. caprinus vs. P. darwini* | 0.000 | 0.000 | 1.00 | 1.00 | 0.0055 | 49.945 |
| **22** | ***P. caprinus vs. P. limatus*** | **0.012** | **0.023** | **0.86** | **0.99** | **0.0055** | **16.690** |
| 23 | *P. caprinus vs. P. magister* | 0.000 | 0.000 | 1.00 | 1.00 | 0.0055 | 29.885 |
| 24 | *P. caprinus vs. P. pehuenche* | 0.000 | 0.000 | 1.00 | 1.00 | 0.0055 | 198.728 |
| 25 | *P. caprinus vs. P. posticalis* | 0.001 | 0.002 | 0.99 | 1.00 | 0.0055 | 70.814 |
| **26** | ***P. caprinus vs. P. vaccarum*** | **0.000** | **0.000** | **1.00** | **1.00** | **0.0055** | **38.060** |
| 27 | *P. caprinus vs. P. xanthopygus* | 0.000 | 0.000 | 1.00 | 1.00 | 0.0055 | 152.185 |
| 28 | *P. chilensis vs. P. darwini* | 0.000 | 0.000 | 1.00 | 1.00 | 0.0055 | 99.982 |
| 29 | *P. chilensis vs. P. limatus* | 0.220 | 0.361 | 0.57 | 0.69 | 0.2640 | 5.644 |
| 30 | *P. chilensis vs. P. magister* | 0.140 | 0.245 | 0.82 | 0.63 | 0.0055 | 20.930 |
| 31 | *P. chilensis vs. P. pehuenche* | 0.000 | 0.000 | 1.00 | 1.00 | 0.0055 | 155.560 |
| **32** | ***P. chilensis vs. P. posticalis*** | **0.073** | **0.137** | **0.83** | **0.89** | **0.0055** | **46.482** |
| 33 | *P. chilensis vs. P. vaccarum* | 0.005 | 0.011 | 0.98 | 0.99 | 0.0055 | 46.990 |
| 34 | *P. chilensis vs. P. xanthopygus* | 0.000 | 0.000 | 1.00 | 1.00 | 0.0055 | 167.217 |
| 35 | *P. darwini vs. P. limatus* | 0.000 | 0.000 | 1.00 | 1.00 | 0.0055 | 81.982 |
| **36** | ***P. darwini vs. P. magister*** | **0.000** | **0.000** | **1.00** | **1.00** | **0.0055** | **82.917** |
| 37 | *P. darwini vs. P. pehuenche* | 0.000 | 0.000 | 1.00 | 1.00 | 0.0055 | 53.887 |
| 38 | *P. darwini vs. P. posticalis* | 0.000 | 0.000 | 1.00 | 1.00 | 0.0055 | 142.201 |
| 39 | *P. darwini vs. P. vaccarum* | 0.103 | 0.188 | 0.65 | 0.87 | 0.0055 | 70.031 |
| 40 | *P. darwini vs. P. xanthopygus* | 0.000 | 0.000 | 1.00 | 1.00 | 0.0055 | 56.678 |
| 41 | *P. limatus vs. P. magister* | 0.331 | 0.498 | 0.66 | 0.06 | 0.7975 | 4.618 |
| 42 | *P. limatus vs. P. pehuenche* | 0.000 | 0.000 | 1.00 | 1.00 | 0.0055 | 104.347 |
| 43 | *P. limatus vs. P. posticalis* | 0.068 | 0.127 | 0.87 | 0.88 | 0.0055 | 47.018 |
| **44** | ***P. limatus vs. P. vaccarum*** | **0.014** | **0.027** | **0.96** | **0.98** | **0.0055** | **46.742** |
| 45 | *P. limatus vs. P. xanthopygus* | 0.000 | 0.000 | 1.00 | 1.00 | 0.0055 | 122.619 |
| 46 | *P. magister vs. P. pehuenche* | 0.000 | 0.000 | 1.00 | 1.00 | 0.0055 | 188.911 |
| 47 | *P. magister vs. P. posticalis* | 0.001 | 0.002 | 1.00 | 1.00 | 0.0055 | 89.035 |
| 48 | *P. magister vs. P. vaccarum* | 0.005 | 0.010 | 0.97 | 0.99 | 0.0055 | 57.022 |
| 49 | *P. magister vs. P. xanthopygus* | 0.000 | 0.000 | 1.00 | 1.00 | 0.0055 | 183.329 |
| 50 | *P. pehuenche vs. P. posticalis* | 0.000 | 0.000 | 1.00 | 1.00 | 0.0055 | 210.288 |
| 51 | *P. pehuenche vs. P. vaccarum* | 0.005 | 0.010 | 0.53 | 1.00 | 0.0055 | 45.355 |
| **52** | ***P. pehuenche vs. P. xanthopygus*** | **0.018** | **0.035** | **0.80** | **0.98** | **0.0055** | **15.195** |
| 53 | *P. posticalis vs. P. vaccarum* | 0.000 | 0.000 | 1.00 | 1.00 | 0.0055 | 129.828 |
| 54 | *P. posticalis vs. P. xanthopygus* | 0.000 | 0.000 | 1.00 | 1.00 | 0.0055 | 223.027 |
| 55 | *P. vaccarum vs. P. xanthopygus* | 0.020 | 0.040 | 0.98 | 0.80 | 0.0055 | 50.102 |

***Results of pairwise comparisons of the fundamental climatic niches of* Phyllotis *species within the* darwini *species group***

**Table S4.** Pairwise comparisons of the fundamental climatic niches of the 11 species of *Phyllotis* within the *darwini* species group. Results of the 55 pairwise comparisons completed using the niche overlap test (NOT) and the niche divergence test (NDT) are depicted. Each case of niche comparison was classified as niche divergence (ND), niche equivalence (NE), or as niche non-equivalence (NNE), following the methodology proposed by Brown & Carnaval (2019) for the joint interpretation of *p*-values. In addition, comparison cases also were categorized into the five discrete intervals of niche overlap described Rödder & Engler (2011), as: no overlap (0-0.2), low overlap (0.2-0.4), moderate overlap (0.4-0.6), high overlap (0.6-0.8), and very high overlap (0.8-1.0), according to the values of the Schoener’s D index obtained. Niche similarity is quantified based on Schoener’s D index of niche overlap, and *p*-values of statistical significance are provided. The percentage of explained variance (% EV) accounted by the two PCs used to perform each pairwise comparison is also shown. Comparisons completed between sister species are those represented in bolded font.

| Case | Species compared | NOT (Full Extension) | | | | NDT (Shared Extension) | | | | % EV | | Case classifications | |
| --- | --- | --- | --- | --- | --- | --- | --- | --- | --- | --- | --- | --- | --- |
|  |  | Niche Similarity | *p*-value | Background 2→1 | Background 1→2 | Niche Similarity | *p*-value | Background 2→1 | Background 1→2 | PC 1 | PC 2 | B & C (2019) | R & E (2011) |
| **1** | ***P. bonariensis vs. P. camiari*** | **0.0000** | **0.0043** | **0.0099** | **0.0099** | **0.0000** | **0.0105** | **0.0099** | **0.0099** | **52.13** | **24.87** | **ND** | **no overlap** |
| 2 | *P. bonariensis vs. P. caprinus* | 0.0000 | 0.0303 | 0.0099 | 0.0099 | 0.0000 | 0.0406 | 0.0099 | 0.0099 | 56.14 | 24.92 | ND | no overlap |
| 3 | *P. bonariensis vs. P. chilensis* | 0.0000 | 0.0099 | 0.0099 | 0.0099 | 0.0000 | 0.0099 | 0.0099 | 0.0099 | 62.98 | 18.62 | ND | no overlap |
| 4 | *P. bonariensis vs. P. darwini* | 0.0000 | 0.0203 | 0.0099 | 0.0099 | 0.0000 | 0.0109 | 0.0099 | 0.0099 | 42.80 | 28.27 | ND | no overlap |
| 5 | *P. bonariensis vs. P. limatus* | 0.0000 | 0.0099 | 0.0100 | 0.0099 | 0.0000 | 0.0099 | 0.0101 | 0.0099 | 57.59 | 23.74 | ND | no overlap |
| 6 | *P. bonariensis vs. P. magister* | 0.0000 | 0.0208 | 0.0100 | 0.0099 | 0.0000 | 0.0102 | 0.0102 | 0.0099 | 56.61 | 27.97 | ND | no overlap |
| 7 | *P. bonariensis vs. P. pehuenche* | 0.0000 | 0.0337 | 0.0099 | 0.0099 | 0.0000 | 0.0307 | 0.0099 | 0.0099 | 52.96 | 30.85 | ND | no overlap |
| 8 | *P. bonariensis vs. P. posticalis* | 0.0000 | 0.0099 | 0.0099 | 0.0099 | 0.0000 | 0.0099 | 0.0099 | 0.0099 | 54.19 | 27.09 | ND | no overlap |
| 9 | *P. bonariensis vs. P. vaccarum* | 0.0000 | 0.0099 | 0.0099 | 0.0099 | 0.0000 | 0.0099 | 0.0099 | 0.0099 | 46.45 | 24.68 | ND | no overlap |
| 10 | *P. bonariensis vs. P. xanthopygus* | 0.0080 | 0.0099 | 0.0081 | 0.0052 | 0.0080 | 0.0099 | 0.0081 | 0.0234 | 46.53 | 30.78 | ND | no overlap |
| 11 | *P. camiari vs. P. caprinus* | 0.0000 | 0.0099 | 0.0099 | 0.0099 | 0.0000 | 0.0099 | 0.0099 | 0.0099 | 56.69 | 23.71 | ND | no overlap |
| 12 | *P. camiari vs. P. chilensis* | 0.0020 | 0.0099 | 0.0436 | 0.0436 | 0.0020 | 0.0099 | 0.0356 | 0.0356 | 58.15 | 24.26 | ND | no overlap |
| 13 | *P. camiari vs. P. darwini* | 0.0130 | NA | 0.0396 | 0.0099 | 0.0130 | 0.0099 | 0.0030 | 0.0099 | 47.16 | 32.08 | ND | no overlap |
| 14 | *P. camiari vs. P. limatus* | 0.0000 | 0.0198 | 0.0101 | 0.0099 | 0.0000 | NA | 0.0099 | 0.0099 | 49.94 | 25.36 | ND | no overlap |
| 15 | *P. camiari vs. P. magister* | 0.0000 | 0.0099 | 0.0103 | 0.0099 | 0.0000 | 0.0099 | 0.0103 | 0.0099 | 50.39 | 28.37 | ND | no overlap |
| 16 | *P. camiari vs. P. pehuenche* | 0.0000 | 0.3267 | 0.0099 | 0.0099 | 0.0000 | 0.2970 | 0.0099 | 0.0099 | 49.95 | 27.01 | ND | no overlap |
| 17 | *P. camiari vs. P. posticalis* | 0.0000 | NA | 0.0099 | 0.0099 | 0.0000 | NA | 0.0099 | 0.0099 | 52.16 | 28.52 | ND | no overlap |
| 18 | *P. camiari vs. P. vaccarum* | 0.0250 | 0.0069 | 0.0406 | 0.0099 | 0.0360 | 0.5743 | 0.0047 | 0.0099 | 47.12 | 28.46 | NNE | no overlap |
| 19 | *P. camiari vs. P. xanthopygus* | 0.0430 | 0.0212 | 0.0163 | 0.0099 | 0.0430 | 0.1980 | 0.0891 | 0.0899 | 53.63 | 30.30 | NNE | no overlap |
| 20 | *P. caprinus vs. P. chilensis* | 0.1710 | 0.3000 | 0.6238 | 0.5050 | 0.1710 | 0.0099 | 0.6832 | 0.7030 | 63.84 | 17.47 | ND | no overlap |
| 21 | *P. caprinus vs. P. darwini* | 0.1190 | 0.0287 | 0.4852 | 0.6437 | 0.1190 | 0.0891 | 0.6238 | 0.5248 | 44.93 | 31.63 | NNE | no overlap |
| **22** | ***P. caprinus vs. P. limatus*** | **0.2130** | **0.0453** | **0.0287** | **0.0292** | **0.2160** | **0.0573** | **0.0213** | **0.0233** | **50.82** | **23.78** | **NNE** | **low overlap** |
| 23 | *P. caprinus vs. P. magister* | 0.2530 | 0.0458 | 0.0451 | 0.0465 | 0.2740 | 1.0000 | 0.0407 | 0.0427 | 48.58 | 25.34 | NNE | low overlap |
| 24 | *P. caprinus vs. P. pehuenche* | 0.0000 | 0.0305 | 0.0099 | 0.0099 | 0.0000 | 0.0011 | 0.0099 | 0.0099 | 37.99 | 35.00 | ND | no overlap |
| 25 | *P. caprinus vs. P. posticalis* | 0.0630 | 0.0099 | 0.8020 | 0.6634 | 0.0630 | 0.0099 | 0.8812 | 0.3663 | 61.30 | 19.84 | ND | no overlap |
| **26** | ***P. caprinus vs. P. vaccarum*** | **0.0740** | **0.0099** | **0.0594** | **0.0297** | **0.0710** | **0.0099** | **0.0257** | **0.0614** | **49.01** | **29.60** | **ND** | **no overlap** |
| 27 | *P. caprinus vs. P. xanthopygus* | 0.0000 | 0.0307 | 0.0099 | 0.0099 | 0.0000 | 0.0406 | 0.0399 | 0.0399 | 40.75 | 35.80 | ND | no overlap |
| 28 | *P. chilensis vs. P. darwini* | 0.1610 | 0.0371 | 0.0415 | 0.0485 | 0.1610 | 0.9901 | 0.4059 | 0.4257 | 47.91 | 25.48 | NNE | no overlap |
| 29 | *P. chilensis vs. P. limatus* | 0.4610 | 0.8812 | 0.0909 | 0.0099 | 0.4600 | 0.8614 | 0.2277 | 0.0099 | 52.55 | 23.95 | NE | moderate overlap |
| 30 | *P. chilensis vs. P. magister* | 0.2830 | 0.0199 | 0.5758 | 0.6238 | 0.2860 | 0.0099 | 0.4592 | 0.4654 | 49.83 | 26.07 | ND | low overlap |
| 31 | *P. chilensis vs. P. pehuenche* | 0.0000 | 0.0109 | 0.0099 | 0.0099 | 0.0000 | 0.0111 | 0.0099 | 0.0099 | 41.53 | 29.80 | ND | no overlap |
| **32** | ***P. chilensis vs. P. posticalis*** | **0.4110** | **0.0289** | **0.0491** | **0.0491** | **0.4110** | **0.0871** | **0.0491** | **0.0495** | **64.26** | **15.42** | **NNE** | **moderate overlap** |
| 33 | *P. chilensis vs. P. vaccarum* | 0.4021 | 0.0375 | 0.0499 | 0.0495 | 0.4021 | 0.0744 | 0.0099 | 0.0099 | 45.02 | 29.16 | NNE | moderate overlap |
| 34 | *P. chilensis vs. P. xanthopygus* | 0.0000 | 0.0403 | 0.0099 | 0.0099 | 0.0000 | 0.0406 | 0.0099 | 0.0099 | 40.79 | 32.37 | ND | no overlap |
| 35 | *P. darwini vs. P. limatus* | 0.0740 | 0.0079 | 0.8700 | 0.6238 | 0.0750 | 0.0099 | 0.9208 | 0.5050 | 45.58 | 36.68 | ND | no overlap |
| **36** | ***P. darwini vs. P. magister*** | **0.0300** | **0.0089** | **0.0478** | **0.0472** | **0.0310** | **0.0099** | **0.0432** | **0.0212** | **47.70** | **26.81** | **ND** | **no overlap** |
| 37 | *P. darwini vs. P. pehuenche* | 0.0010 | 0.0099 | 0.9406 | 0.9802 | 0.0010 | 0.0099 | 0.9208 | 0.9010 | 45.69 | 29.84 | ND | no overlap |
| 38 | *P. darwini vs. P. posticalis* | 0.0850 | 0.0494 | 0.0436 | 0.0822 | 0.0840 | 0.0494 | 0.0454 | 0.0582 | 42.68 | 27.40 | ND | no overlap |
| 39 | *P. darwini vs. P. vaccarum* | 0.3020 | 0.0413 | 0.0475 | 0.0396 | 0.3020 | 0.0832 | 0.0277 | 0.0238 | 45.61 | 29.39 | NNE | low overlap |
| 40 | *P. darwini vs. P. xanthopygus* | 0.0260 | 0.0099 | 0.0287 | 0.0139 | 0.0260 | 0.0099 | 0.0455 | 0.0416 | 44.70 | 27.40 | ND | no overlap |
| 41 | *P. limatus vs. P. magister* | 0.5140 | 0.0399 | 0.0909 | 0.1485 | 0.4940 | 0.0099 | 0.0153 | 0.1742 | 41.89 | 38.05 | ND | moderate overlap |
| 42 | *P. limatus vs. P. pehuenche* | 0.0280 | 0.0198 | 0.0218 | 0.0321 | 0.0280 | 0.0198 | 0.0337 | 0.0139 | 46.75 | 27.27 | ND | no overlap |
| 43 | *P. limatus vs. P. posticalis* | 0.2850 | 0.0402 | 0.0594 | 0.0238 | 0.2860 | 0.0801 | 0.0644 | 0.6040 | 57.56 | 21.59 | NNE | low overlap |
| **44** | ***P. limatus vs. P. vaccarum*** | **0.3490** | **0.0241** | **0.0589** | **0.0589** | **0.3330** | **0.0576** | **0.0493** | **0.0411** | **49.98** | **26.14** | **NNE** | **low overlap** |
| 45 | *P. limatus vs. P. xanthopygus* | 0.0000 | 0.0099 | 0.0099 | 0.0099 | 0.0000 | 0.0099 | 0.0099 | 0.0099 | 47.09 | 26.91 | ND | no overlap |
| 46 | *P. magister vs. P. pehuenche* | 0.0000 | 0.0099 | 0.0099 | 0.0099 | 0.0000 | 0.0099 | 0.0099 | 0.0103 | 46.40 | 27.11 | ND | no overlap |
| 47 | *P. magister vs. P. posticalis* | 0.2350 | 0.2871 | 0.4455 | 0.9286 | 0.2320 | 0.8713 | 0.3861 | 0.5446 | 55.50 | 23.57 | NE | low overlap |
| 48 | *P. magister vs. P. vaccarum* | 0.2160 | 0.0942 | 0.6238 | 0.6020 | 0.2090 | 0.9872 | 0.6832 | 0.5258 | 41.27 | 29.82 | NE | low overlap |
| 49 | *P. magister vs. P. xanthopygus* | 0.0000 | 0.0495 | 0.0099 | 0.0101 | 0.0000 | 0.0099 | 0.0099 | 0.0099 | 42.62 | 28.84 | ND | no overlap |
| 50 | *P. pehuenche vs. P. posticalis* | 0.0000 | 0.0483 | 0.0099 | 0.0099 | 0.0000 | 0.1485 | 0.0099 | 0.0099 | 44.34 | 25.72 | NNE | no overlap |
| 51 | *P. pehuenche vs. P. vaccarum* | 0.3100 | 0.0982 | 0.0693 | 0.3069 | 0.3070 | 0.9703 | 0.0693 | 0.2079 | 44.49 | 27.85 | NE | low overlap |
| **52** | ***P. pehuenche vs. P. xanthopygus*** | **0.3770** | **0.0481** | **0.2673** | **0.0663** | **0.3770** | **0.0921** | **0.0911** | **0.0861** | **46.84** | **23.79** | **NNE** | **low overlap** |
| 53 | *P. posticalis vs. P. vaccarum* | 0.2850 | 0.0471 | 0.0189 | 0.0089 | 0.2850 | 0.0372 | 0.0128 | 0.0393 | 45.37 | 28.46 | ND | low overlap |
| 54 | *P. posticalis vs. P. xanthopygus* | 0.0000 | 0.0287 | 0.0099 | 0.0099 | 0.0000 | 0.0172 | 0.0099 | 0.0099 | 42.65 | 32.26 | ND | no overlap |
| 55 | *P. vaccarum vs. P. xanthopygus* | 0.1450 | 0.0278 | 0.0049 | 0.0454 | 0.2860 | 0.0374 | 0.0463 | 0.0396 | 44.61 | 29.68 | ND | no overlap |

***Results of pairwise quantifications of niche overlap between the species of* Phyllotis *within the* darwini *species group across the selected bioclimatic variables***

**Table S5.** Quantifications of climatic niche overlaps among 11 species of *Phyllotis* across the selected bioclimatic variables. Quantifications of niche overlap are based on Schoener’s D metric and were completed based on the profiles of predicted niche occupancy (PNO). Values of Schoener’s D are depicted for the 55 resulting pairwise comparisons, and those corresponding to sister species are represented in bold.

| Case | Species compared | Bio 02 | Bio 04 | Bio 06 | Bio 08 | Bio 13 | Bio 14 | Bio 15 | Bio 19 | Average of niche overlap  per pairwise comparison |
| --- | --- | --- | --- | --- | --- | --- | --- | --- | --- | --- |
| **1** | ***P. bonariensis vs. P. camiari*** | **0.873** | **0.819** | **0.898** | **0.751** | **0.914** | **0.571** | **0.659** | **0.597** | **0.760** |
| 2 | *P. bonariensis vs. P. caprinus* | 0.678 | 0.252 | 0.539 | 0.452 | 0.704 | 0.062 | 0.175 | 0.185 | 0.381 |
| 3 | *P. bonariensis vs. P. chilensis* | 0.438 | 0.107 | 0.362 | 0.314 | 0.532 | 0.365 | 0.326 | 0.370 | 0.352 |
| 4 | *P. bonariensis vs. P. darwini* | 0.574 | 0.203 | 0.681 | 0.144 | 0.388 | 0.382 | 0.361 | 0.404 | 0.392 |
| 5 | *P. bonariensis vs. P. limatus* | 0.475 | 0.055 | 0.309 | 0.374 | 0.546 | 0.265 | 0.355 | 0.336 | 0.339 |
| 6 | *P. bonariensis vs. P. magister* | 0.492 | 0.041 | 0.434 | 0.332 | 0.507 | 0.010 | 0.272 | 0.029 | 0.265 |
| 7 | *P. bonariensis vs. P. pehuenche* | 0.684 | 0.825 | 0.273 | 0.053 | 0.530 | 0.358 | 0.747 | 0.571 | 0.505 |
| 8 | *P. bonariensis vs. P. posticalis* | 0.602 | 0.010 | 0.297 | 0.198 | 0.481 | 0.352 | 0.225 | 0.640 | 0.351 |
| 9 | *P. bonariensis vs. P. vaccarum* | 0.561 | 0.550 | 0.483 | 0.447 | 0.350 | 0.258 | 0.571 | 0.421 | 0.455 |
| 10 | *P. bonariensis vs. P. xanthopygus* | 0.605 | 0.722 | 0.359 | 0.190 | 0.398 | 0.437 | 0.734 | 0.618 | 0.508 |
| 11 | *P. camiari vs. P. caprinus* | 0.849 | 0.494 | 0.769 | 0.714 | 0.897 | 0.534 | 0.530 | 0.668 | 0.682 |
| 12 | *P. camiari vs. P. chilensis* | 0.647 | 0.260 | 0.531 | 0.606 | 0.717 | 0.691 | 0.668 | 0.637 | 0.595 |
| 13 | *P. camiari vs. P. darwini* | 0.668 | 0.469 | 0.894 | 0.304 | 0.604 | 0.722 | 0.691 | 0.614 | 0.621 |
| 14 | *P. camiari vs. P. limatus* | 0.677 | 0.193 | 0.494 | 0.624 | 0.759 | 0.633 | 0.653 | 0.629 | 0.583 |
| 15 | *P. camiari vs. P. magister* | 0.724 | 0.158 | 0.651 | 0.431 | 0.739 | 0.261 | 0.427 | 0.298 | 0.461 |
| 16 | *P. camiari vs. P. pehuenche* | 0.893 | 0.947 | 0.400 | 0.176 | 0.770 | 0.942 | 0.963 | 0.535 | 0.703 |
| 17 | *P. camiari vs. P. posticalis* | 0.751 | 0.087 | 0.426 | 0.611 | 0.504 | 0.911 | 0.760 | 0.794 | 0.605 |
| 18 | *P. camiari vs. P. vaccarum* | 0.796 | 0.795 | 0.669 | 0.683 | 0.624 | 0.794 | 0.796 | 0.794 | 0.744 |
| 19 | *P. camiari vs. P. xanthopygus* | 0.778 | 0.938 | 0.471 | 0.314 | 0.649 | 0.914 | 0.793 | 0.754 | 0.701 |
| 20 | *P. caprinus vs. P. chilensis* | 0.908 | 0.899 | 0.843 | 0.917 | 0.900 | 0.900 | 0.917 | 0.906 | 0.899 |
| 21 | *P. caprinus vs. P. darwini* | 0.673 | 0.912 | 0.937 | 0.672 | 0.837 | 0.900 | 0.933 | 0.618 | 0.810 |
| **22** | ***P. caprinus vs. P. limatus*** | **0.907** | **0.853** | **0.834** | **0.905** | **0.933** | **0.940** | **0.866** | **0.934** | **0.897** |
| 23 | *P. caprinus vs. P. magister* | 0.948 | 0.791 | 0.920 | 0.808 | 0.899 | 0.869 | 0.784 | 0.853 | 0.859 |
| 24 | *P. caprinus vs. P. pehuenche* | 0.926 | 0.485 | 0.659 | 0.220 | 0.930 | 0.501 | 0.534 | 0.282 | 0.567 |
| 25 | *P. caprinus vs. P. posticalis* | 0.936 | 0.476 | 0.709 | 0.775 | 0.597 | 0.762 | 0.818 | 0.726 | 0.725 |
| **26** | ***P. caprinus vs. P. vaccarum*** | **0.954** | **0.801** | **0.899** | **0.872** | **0.833** | **0.881** | **0.786** | **0.858** | **0.861** |
| 27 | *P. caprinus vs. P. xanthopygus* | 0.787 | 0.593 | 0.659 | 0.462 | 0.824 | 0.516 | 0.427 | 0.381 | 0.581 |
| 28 | *P. chilensis vs. P. darwini* | 0.664 | 0.740 | 0.711 | 0.838 | 0.925 | 0.981 | 0.918 | 0.759 | 0.817 |
| 29 | *P. chilensis vs. P. limatus* | 0.975 | 0.957 | 0.937 | 0.949 | 0.921 | 0.969 | 0.972 | 0.970 | 0.956 |
| 30 | *P. chilensis vs. P. magister* | 0.963 | 0.923 | 0.885 | 0.892 | 0.808 | 0.811 | 0.888 | 0.815 | 0.873 |
| 31 | *P. chilensis vs. P. pehuenche* | 0.804 | 0.250 | 0.644 | 0.404 | 0.897 | 0.636 | 0.668 | 0.551 | 0.607 |
| **32** | ***P. chilensis vs. P. posticalis*** | **0.928** | **0.726** | **0.929** | **0.895** | **0.785** | **0.831** | **0.839** | **0.852** | **0.848** |
| 33 | *P. chilensis vs. P. vaccarum* | 0.945 | 0.587 | 0.866 | 0.887 | 0.833 | 0.934 | 0.873 | 0.930 | 0.857 |
| 34 | *P. chilensis vs. P. xanthopygus* | 0.764 | 0.311 | 0.602 | 0.637 | 0.852 | 0.700 | 0.535 | 0.578 | 0.622 |
| 35 | *P. darwini vs. P. limatus* | 0.762 | 0.692 | 0.713 | 0.783 | 0.942 | 0.988 | 0.875 | 0.652 | 0.801 |
| **36** | ***P. darwini vs. P. magister*** | **0.640** | **0.607** | **0.848** | **0.883** | **0.864** | **0.838** | **0.730** | **0.427** | **0.730** |
| 37 | *P. darwini vs. P. pehuenche* | 0.781 | 0.474 | 0.569 | 0.605 | 0.809 | 0.654 | 0.724 | 0.812 | 0.679 |
| 38 | *P. darwini vs. P. posticalis* | 0.800 | 0.229 | 0.571 | 0.665 | 0.677 | 0.827 | 0.868 | 0.725 | 0.670 |
| 39 | *P. darwini vs. P. vaccarum* | 0.765 | 0.816 | 0.829 | 0.831 | 0.861 | 0.947 | 0.890 | 0.860 | 0.850 |
| 40 | *P. darwini vs. P. xanthopygus* | 0.954 | 0.617 | 0.591 | 0.784 | 0.796 | 0.692 | 0.617 | 0.809 | 0.732 |
| 41 | *P. limatus vs. P. magister* | 0.968 | 0.987 | 0.896 | 0.888 | 0.950 | 0.904 | 0.946 | 0.897 | 0.930 |
| 42 | *P. limatus vs. P. pehuenche* | 0.826 | 0.176 | 0.539 | 0.478 | 0.865 | 0.575 | 0.667 | 0.395 | 0.565 |
| 43 | *P. limatus vs. P. posticalis* | 0.924 | 0.659 | 0.871 | 0.829 | 0.635 | 0.776 | 0.796 | 0.781 | 0.784 |
| **44** | ***P. limatus vs. P. vaccarum*** | **0.967** | **0.529** | **0.836** | **0.902** | **0.881** | **0.932** | **0.884** | **0.898** | **0.854** |
| 45 | *P. limatus vs. P. xanthopygus* | 0.835 | 0.199 | 0.503 | 0.691 | 0.821 | 0.605 | 0.555 | 0.473 | 0.585 |
| 46 | *P. magister vs. P. pehuenche* | 0.850 | 0.140 | 0.653 | 0.478 | 0.850 | 0.220 | 0.461 | 0.097 | 0.469 |
| 47 | *P. magister vs. P. posticalis* | 0.900 | 0.672 | 0.693 | 0.693 | 0.424 | 0.458 | 0.599 | 0.446 | 0.610 |
| 48 | *P. magister vs. P. vaccarum* | 0.976 | 0.459 | 0.933 | 0.856 | 0.930 | 0.738 | 0.772 | 0.684 | 0.793 |
| 49 | *P. magister vs. P. xanthopygus* | 0.766 | 0.135 | 0.624 | 0.730 | 0.831 | 0.246 | 0.423 | 0.131 | 0.486 |
| 50 | *P. pehuenche vs. P. posticalis* | 0.903 | 0.097 | 0.556 | 0.408 | 0.500 | 0.920 | 0.738 | 0.641 | 0.595 |
| 51 | *P. pehuenche vs. P. vaccarum* | 0.927 | 0.705 | 0.814 | 0.596 | 0.905 | 0.809 | 0.857 | 0.684 | 0.787 |
| **52** | ***P. pehuenche vs. P. xanthopygus*** | **0.898** | **0.948** | **0.967** | **0.945** | **0.944** | **0.965** | **0.885** | **0.934** | **0.936** |
| 53 | *P. posticalis vs. P. vaccarum* | 0.935 | 0.137 | 0.724 | 0.759 | 0.377 | 0.925 | 0.762 | 0.828 | 0.681 |
| 54 | *P. posticalis vs. P. xanthopygus* | 0.862 | 0.053 | 0.533 | 0.570 | 0.423 | 0.933 | 0.499 | 0.699 | 0.572 |
| 55 | *P. vaccarum vs. P. xanthopygus* | 0.878 | 0.823 | 0.831 | 0.774 | 0.943 | 0.822 | 0.862 | 0.773 | 0.838 |
|  | Average of niche overlap  per climatic dimensions | 0.799 | 0.516 | 0.674 | 0.618 | 0.738 | 0.679 | 0.689 | 0.628 |  |

***Spatial projections and performances of the climatic niche models (CNMs) developed for the 11 species members of* Phyllotis darwini *species group***

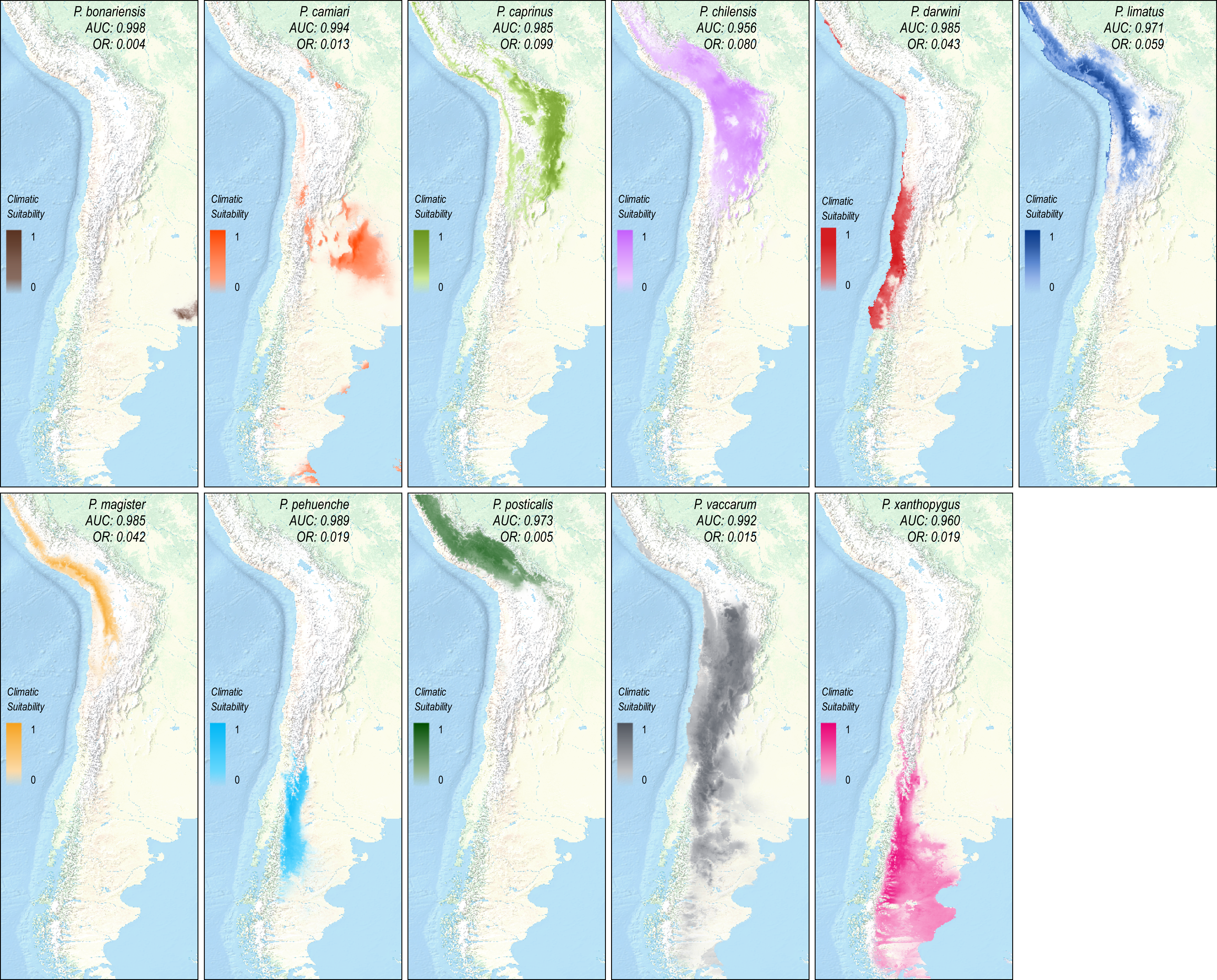

**Figure S2.** Spatial projections and performances of the climatic niche models (CNM) developed for the *Phyllotis* species within the *darwini* species group. Values of the area under the receiver operating characteristic curve test (AUC) and the omission rate under the tenth percentile of training presence (OR) are depicted for each model. Values of climatic suitability are represented as a gradual pattern of coloration where darker tones represent higher values of environmental (climatic) suitability. Projections of these models provided a realistic estimation of the potential geographic distribution of these species, as can be verified by looking the observed geographic distributions of these species in Figure 1.

***Evaluation of the phylogenetic signal in the variations sampled across the bioclimatic variables considered***

The phylogenetic signal was evaluated across the variations sampled in the bioclimatic variables by inputting the median values and the ultrametric tree in the R package *phylosignal* (Keck et al. 2016), running 1000 random subsamples, and employing five different metrics (Abouheif’s Cmean, Moran’s I, Blomberg’s K and K*, as well as Pagel’s λ).

**Table S6.** Evaluation of the phylogenetic signal across the bioclimatic variables considered. The metrics of Phylogenetic signal employed were: Abouheif’s Cmean (C), Moran’s I (I), Blomberg’s K (K) and K* (K*), as well as Pagel’s λ (λ). The phylogenetic signal evaluated was considered significant when *p*-values obtained were less than 0.05.

| Bioclimatic variable | C | *p*-value | I | *p*-value | K | *p*-value | K* | *p*-value | λ | *p*-value |
| --- | --- | --- | --- | --- | --- | --- | --- | --- | --- | --- |
| Bio 2: Annual mean diurnal range | -0.3490 | 0.8971 | -0.1270 | 0.8312 | 0.7397 | 0.3676 | 0.8163 | 0.2927 | 0.0981 | 1.0000 |
| Bio 4: Temperature seasonality | 0.2904 | 0.2350 | -0.0507 | 0.0529 | 0.9878 | 0.0639 | 1.0144 | 0.0689 | 0.8559 | 0.4849 |
| Bio 6: Min Temperature of coldest month | -0.0465 | 0.3806 | -0.0966 | 0.4286 | 0.7494 | 0.3437 | 0.8228 | 0.3117 | 0.1651 | 1.0000 |
| Bio 8: Mean temperature of wettest quarter | 0.3232 | 0.1250 | -0.0713 | 0.1688 | 0.8354 | 0.1938 | 0.9212 | 0.1389 | 0.0831 | 1.0000 |
| Bio 13: Precipitation of wettest month | 0.2016 | 0.1350 | -0.0553 | 0.0589 | 0.9928 | 0.0889 | 1.0961 | 0.0659 | 1.2135 | 0.2144 |
| Bio 14: Precipitation of driest month | 0.2508 | 0.0720 | -0.0601 | 0.0899 | 0.9072 | 0.1339 | 0.9668 | 0.1249 | 0.9588 | 1.0000 |
| Bio 15: Precipitation seasonality | 0.2463 | 0.0969 | -0.0585 | 0.0839 | 0.9633 | 0.0739 | 0.9605 | 0.0829 | 0.5071 | 0.6962 |
| Bio 19: Precipitation of coldest quarter | -0.1990 | 0.6234 | -0.1204 | 0.7233 | 0.7454 | 0.3646 | 0.8178 | 0.2727 | 0.0754 | 1.0000 |

In addition, simulations were performed to explore the behavior of the phylogenetic signal metrics based on the phylogenetic information provided by the dated phylogenetic tree, under varying amounts of Brownian motion, with 1000 simulated solutions and 99 repetitions for *p*-value estimation. The number of solutions showing high phylogenetic signals was considerably low (Figure S3).

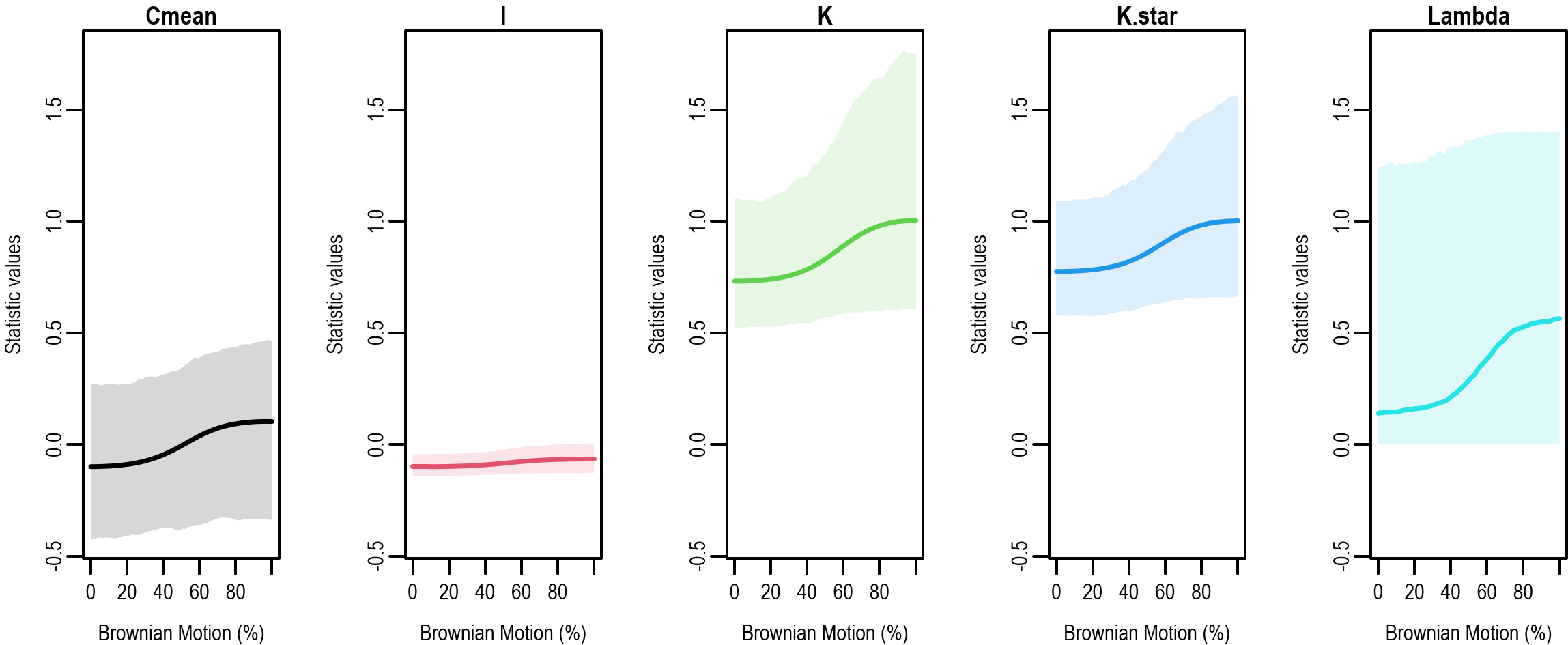

**Figure S3.** Phylogeny-based simulations showing the distribution of values (shades represent 95 % confidence intervals) for each of the five metrics measuring the phylogenetic signal under different amounts of Brownian motion.

The performances of all metrics were very similar in terms of the frequency of significant phylogenetic signals (Figure S4). Up to 50% Brownian motion, the frequency was close to zero, and gradually increased up to about 20% and around 80% Brownian motion, where it reached a plateau.

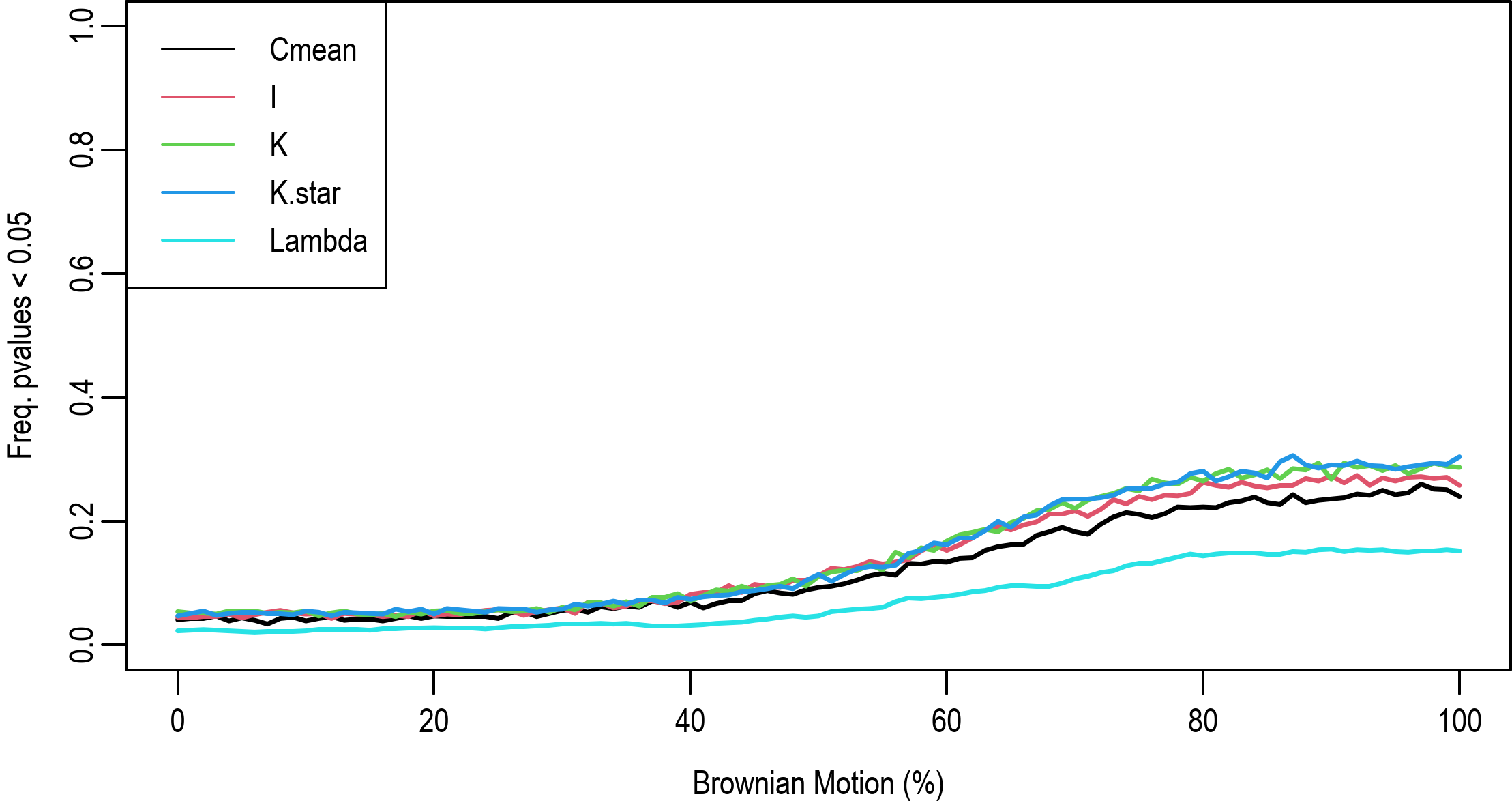

**Figure S4.** Frequency of significant *p*-values (i.e., *p* < 0.05) estimated for each of the five metrics employed to quantifying the phylogenetic signal under varying amounts of Brownian motion.

According to all these results, no significant phylogenetic signal was found (Table S6) and, in general, the values of the metrics for quantifying phylogenetic signal were also low. At the same time, were did not find higher probability of detecting significant phylogenetic signal values along the phylogenetic tree or among the bioclimatic variables considered (Figures S3 and S4).

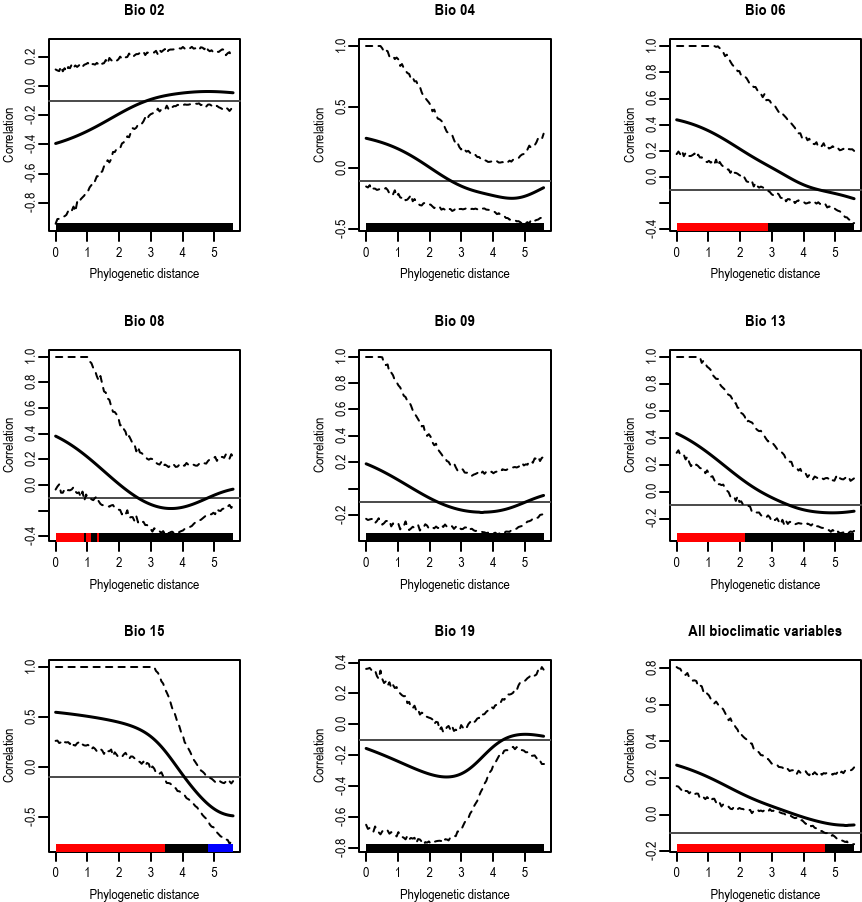

**Figure S5.** Phylogenetic correlograms (Moran’s I) computed for each bioclimatic variable with 95% confidence intervals. Significant positive (red) and/or negative (blue) correlations are marked along the x-axis in each plot.

Statistically significant positive correlations were found in minimum temperature of the coldest month (Bio 6), average temperature of the wettest quarter (Bio 8), precipitation of the wettest month (Bio 13) and seasonality of precipitation (Bio 15), where a significant negative correlation was also found. For the joint evaluation of all bioclimatic variables, the overall correlation was positive and significant.

***Local Indicator of Phylogenetic Association (LIPA)***

The Local Indicator of Phylogenetic Association (LIPA) tests completed using the two-sided alternative hypothesis only shows a statistically significant and negative association between the phylogenetic distances calculated among species and the climatic niches variations in *Phyllotis chilensis* for the annual mean diurnal range (Bio 02), and in *P*. *darwini* for precipitation of coldest quarter (Bio 19).

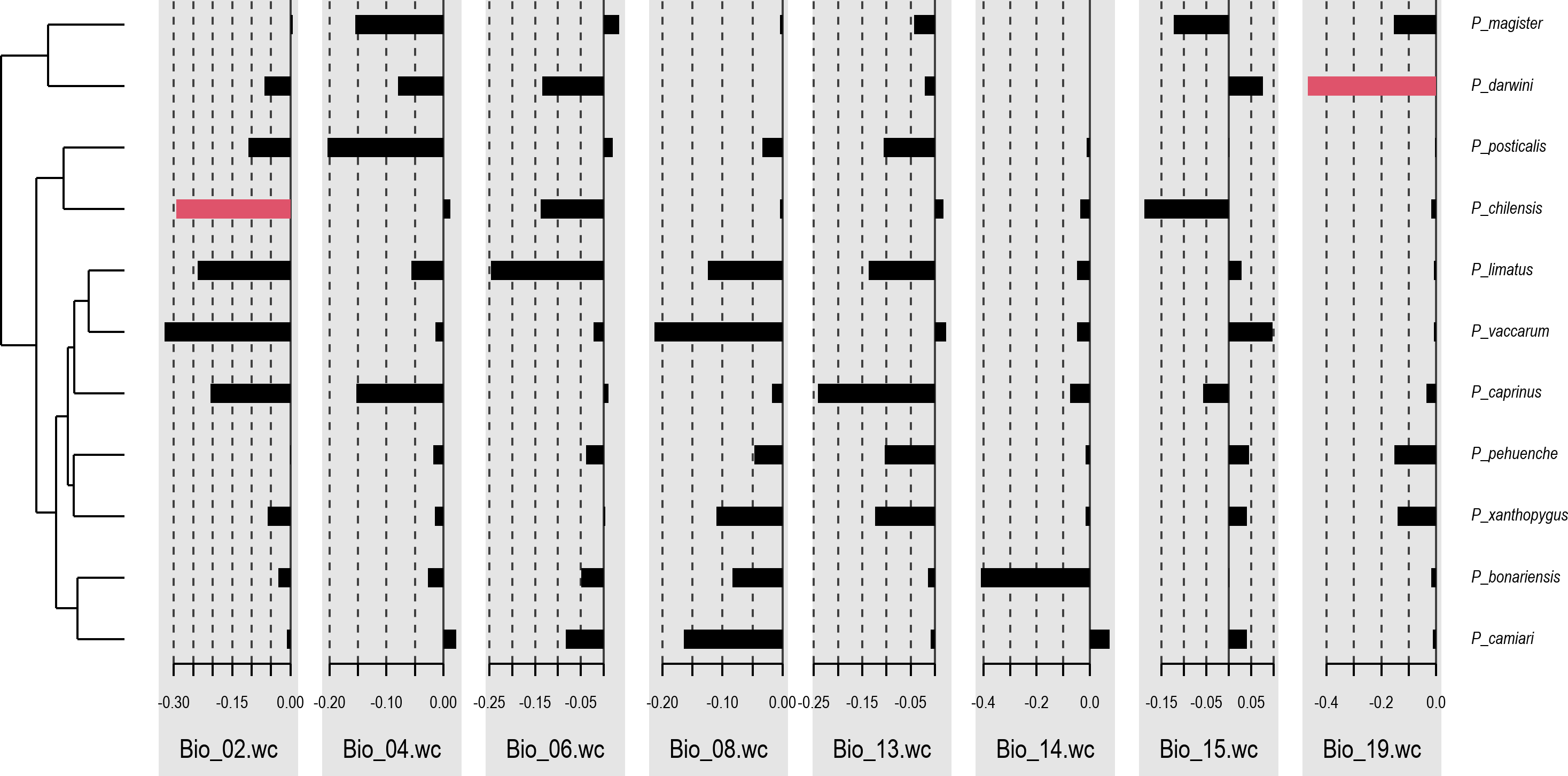

**Figure S6.** Local Indicator of Phylogenetic Association (local Moran’s I) for each bioclimatic variable. Red bars show statically significant values (based on two-sided tests) for the respective species (right). The tree topology is shown on the left for orientation about phylogenetic relationships.
